# Limits to evolutionary rescue by conjugative plasmids

**DOI:** 10.1101/2022.12.07.519465

**Authors:** Félix Geoffroy, Hildegard Uecker

## Abstract

Plasmids may carry genes coding for beneficial traits and thus contribute to adaptation of bacterial populations to environmental stress. Conjugative plasmids can horizontally transfer between cells, which a priori facilitates the spread of adaptive alleles. However, if the potential recipient cell is already colonized by another incompatible plasmid, successful transfer may be prevented. Competition between plasmids can thus limit horizontal transfer. Previous modeling has indeed shown that evolutionary rescue by a conjugative plasmid is hampered by incompatible resident plasmids in the population. If the rescue plasmid is a mutant variant of the resident plasmid, both plasmids transfer at the same rates. A high conjugation rate then has two, potentially opposing, effects – a direct positive effect on spread of the rescue plasmid and an increase in the fraction of resident plasmid cells. This raises the question whether a high conjugation rate always benefits evolutionary rescue. In this article, we systematically analyse three models of increasing complexity to disentangle the benefits and limits of increasing horizontal gene transfer in the presence of plasmid competition and plasmid costs. We find that the net effect can be positive or negative and that the optimal transfer rate is thus not always the highest one. These results can contribute to our understanding of the many facets of plasmid-driven adaptation and the wide range of transfer rates observed in nature.

## 1 Introduction

Populations can escape extinction after a sudden environmental change if a well-adapted genotype appears in the population and escapes stochastic loss before the resident population goes extinct. This process is referred to as evolutionary rescue [Gomulkiewicz and Holt, 1995, Orr and Unckless, 2008, Bell, 2017], see Figure 1a. In bacterial populations, genes can be located on the chromosome or on extrachromosomal DNA elements such as plasmids [Thomas, 2000]. While many of the genes on plasmids are devoted to the survival and reproduction of the plasmid, some of them are also beneficial for the bacterial host. Beneficial functions include virulence, heavy metal tolerance, and antibiotic resistance [Rankin et al., 2011, Harrison and Brockhurst, 2012, Carroll and Wong, 2018]. It has been found that in commensal *Escherichia coli*, antibiotic resistance genes are more often located on plasmids than on the bacterial chromosome [Stephens et al., 2020]. Adaptation on plasmids differs from adaptation on the chromosome. The maybe most striking difference is that some types of plasmids can transfer between cells of the same and even between cells of different species, making genes on plasmids more mobile than genes on the chromosome. About one quarter of plasmids are conjugative, i.e. possess all genes required for autonomous transfer [Coluzzi et al., 2022]. In conjugation, a donor cell replicates the plasmid and produces a protein complex which allows the injection of the copy into a recipient cell.

**Figure 1.**
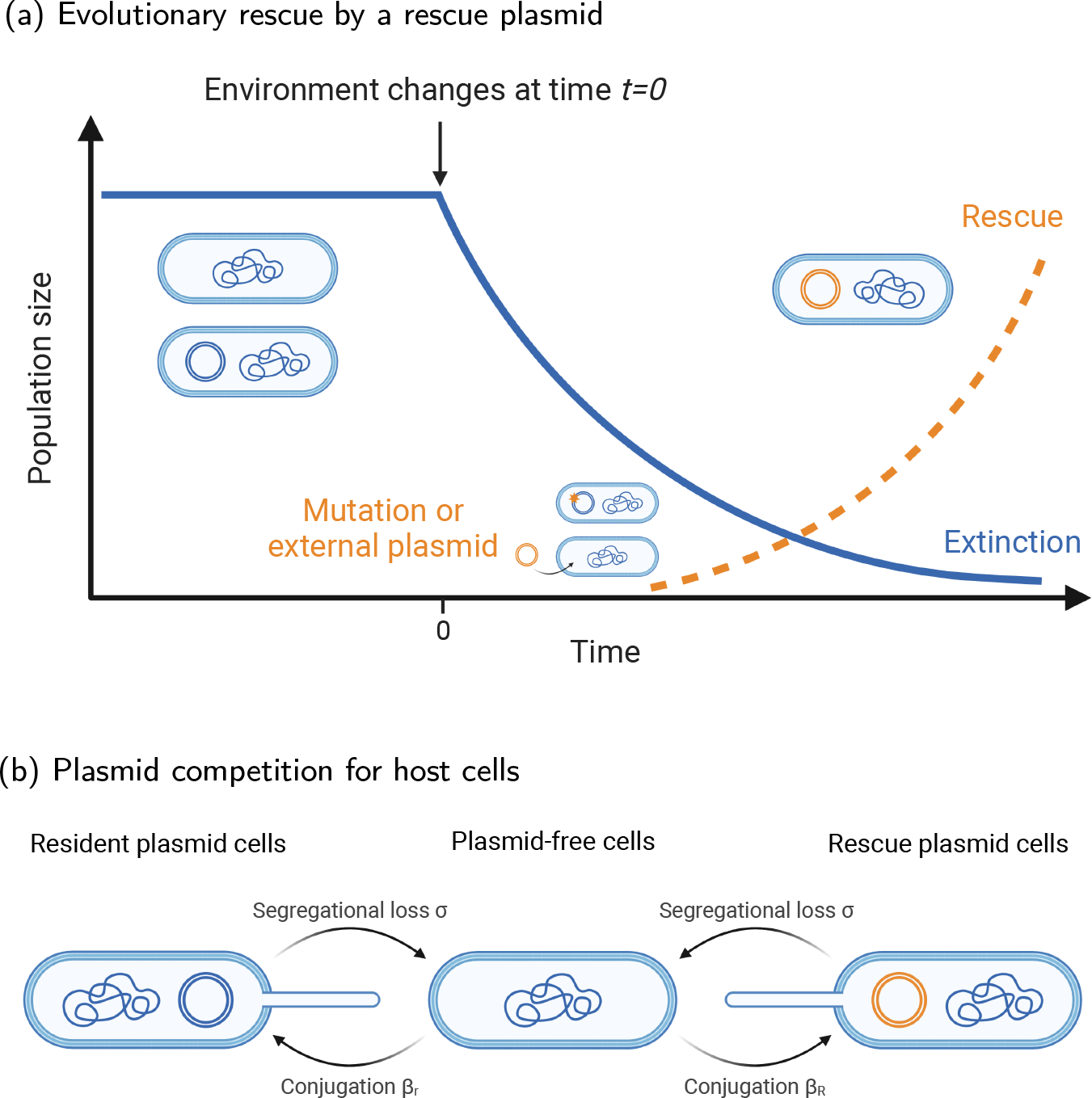
Sketches of **(a)** evolutionary rescue on a conjugative plasmid and **(b)** plasmid competition for hosts. Panel (a) shows the dynamics of a population undergoing evolutionary rescue driven by the appearance and establishment of a beneficial rescue plasmid. Prior to change, the population consists of a mix of plasmid-free cells and cells carrying a resident plasmid (blue). After the change, the population size declines. An adaptive rescue plasmid (orange) might appear either through a mutation on the resident plasmid or through immigration from an external population. Panel (b) illustrates the transitions between the three types of cells (plasmid-free cells, cells carrying the resident plasmid, cells carrying the rescue plasmid). Plasmids can horizontally transfer into plasmid-free cells at rates *β*_*r*_ (resident plasmids) and *β*_*R*_ (rescue plasmids). In the main text, we assume that *β*_*r*_ = *β*_*R*_. In SI section S3, we study rescue if the transfer coefficients of the two plasmid types are proportional to each to each other *β*_*R*_ = *γβ*_*r*_ or independent from each other. Segregational loss of plasmids re-generates plasmid-free cells. In the main text, we ignore this possibility. We explore the effects of segregational loss in SI section S4.

Horizontal gene transfer, especially via conjugation, is considered to be an important driver of bacterial adaptation [Frost et al., 2005, Soucy et al., 2015], including the evolution of antibiotic resistance [San Millan, 2018]. In fact, several instances of rescue occurring with the help of a plasmid have been reported [Ojala et al., 2014, Mattila et al., 2017]. Previous theoretical studies have addressed the role of horizontal transfer in bacterial adaptation. Most of them investigate the effect of conjugation on the fixation probability of a beneficial allele [Bichsel et al., 2010, Novozhilov et al., 2005, Tazzyman and Bonhoeffer, 2013, Billiard et al., 2016]. They all conclude that conjugation increases the probability of adaptation, suggesting the higher the transfer rate the better. Consistently, Sheppard et al. [2021] find that the conjugation rate of a plasmid carrying a beneficial gene evolves to higher values, at least if a high transfer rate is not too costly by itself.

Little attention has been paid to the limits of adaptation on conjugative plasmids. One potential important limitation is the competition of the ‘rescue plasmid’ with other plasmids that are already present in the bacterial population (in the following termed ‘resident plasmids’). If another plasmid that uses the same replication or segregation mechanism is already present in the recipient cell, it is possible that the two plasmids cannot stably coexist and one of them is lost at subsequent cell divisions. These plasmids are said to be incompatible [Novick, 1987]. Many plasmids furthermore code for entry or surface exclusion that prevent entry of incompatible plasmids up front [Garcillán-Barcia and de la Cruz, 2008]. Such incompatible plasmids would impair the spread of a new beneficial conjugative plasmid by preventing entry into their host cells.

Plasmid competition has been considered in the specific contexts of the evolution of cooperative genes [Mc Ginty et al., 2011, Dimitriu et al., 2014] and the questions of why and when certain genes are frequently located on plasmids [Svara and Rankin, 2011, Lehtinen et al., 2021]. To our knowledge, a single study – Tazzyman and Bonhoeffer [2014] – addresses the influence of plasmid competition in the context of evolutionary rescue. In their model, the authors assume that a resident plasmid segregates in the population at some given frequency. This resident plasmid provides no benefit to its carrier. The adaptive rescue plasmid may appear through a *de novo* mutation of a gene on the resident plasmid or through immigration from an external population. Tazzyman and Bonhoeffer [2014] conclude that, even though the presence of a large number of resident plasmids increases the chances of appearance of beneficial alleles via *de novo* mutations, prevalent resident plasmids can sometimes lower the probability of rescue because they compete with the rescue plasmids for host cells, reducing both the rate of immigration and the spread of incompatible rescue plasmids.

In the model by Tazzyman and Bonhoeffer [2014], the fraction of cells that carry the resident plasmid is a fixed exogenous parameter that does not change over time as the population heads towards extinction. As noted by the authors, in reality the fraction of cells that carry the resident plasmid is an endogenous property of the plasmid, which depends, among other things, on the rate at which the resident plasmid is transferred and can change with time. Tazzyman and Bonhoeffer [2014] furthermore assume that this fraction is independent from the transfer coefficient of the rescue plasmid. I.e., the fraction of resident plasmids and the transfer coefficient of the rescue plasmid are independent model parameters. Higher transfer coefficients for the rescue plasmid always promote evolutionary rescue since more transfer facilitates its spread, irrespective of the fraction of available potential recipient cells. The study thus comes to the same conclusion regarding the role of conjugation in adaptation as studies that do not include plasmid competition.

Treating the fraction of resident plasmids and the propensity of the rescue plasmid to transfer as independent is reasonable when both plasmids are sufficiently distinct such that their transfer rates are independent from each other. However, when the rescue plasmid is a mutant variant of the resident plasmid, either generated by a *de novo* mutation *in situ* or within another population, we would expect that both transfer at the same rate. Even if the two plasmids are distinct, their transfer rates might still be positively correlated if transfer is (at least partially) under host control. In these cases, the frequency of the resident plasmid and the transfer coefficient of the rescue plasmid would be positively correlated as well. A higher transfer coefficient now becomes a double-edged sword. On the one hand, higher transfer rates have a direct positive effect on the spread of the rescue plasmid. By increasing the fraction of resident plasmids, they moreover make *de novo* mutations on the plasmid more likely. On the other hand, since the fraction of plasmid-free cells is lower, immigration of incompatible plasmids is less likely and moreover has an indirect negative effect on spread of the rescue plasmid. These antagonistic effects of horizontal plasmid transfer raise the question whether more horizontal transfer always increases the probability of rescue.

In this article, to address this question, we extend the model by Tazzyman and Bonhoeffer [2014] and explicitly consider the dynamics of the resident plasmid and thus also the exact dynamics of plasmid competition for host cells, assuming in the main text that both plasmid types transfer at the same rate (Figure 1b). We consider three models of increasing complexity. The first model is (up to a minor modification) equivalent to the model in Tazzyman and Bonhoeffer [2014] and serves for comparison. In the second model, the dynamics of the resident plasmid are explicitly modeled after the harmful environmental change, but the frequency at the time of change is an exogenous parameter. Finally, in the third model, we determine the initial frequency of the resident plasmid by a balance between horizontal transfer and plasmid costs prior to the environmental deterioration. We find that in all three models, when rescue plasmids appear via *de novo* mutations on resident plasmids, an increase in the transfer coefficient promotes rescue unless the metabolic cost of carrying a plasmid is very high. In contrast, when rescue relies on the immigration of a rescue plasmid that is incompatible with a resident plasmid and has the same propensity to transfer, higher transfer coefficients mostly reduce the probability of evolutionary rescue, at least when the plasmid immigration rate is not correlated with the within-population transfer rate. Last, we show that, even when the rescue plasmid is compatible with the resident plasmid, i.e. in the absence of competition at the plasmid-level, horizontal transfer can reduce the probability of rescue if the population dynamics before the environmental change are taken into account.

## 2 Methods

### 2.1 General model

We consider a bacterial population in a volume *V* that is exposed to a sudden change in its environment at time *t* = 0. We distinguish three types of cells: plasmid-free cells (with population size *N*_*F*_), cells carrying a resident plasmid (*N*_*r*_), and cells carrying a rescue plasmid (*N*_*R*_). We call the population of cells that are either plasmid-free or carry the resident plasmid, (*N*_*F*_ + *N*_*r*_), the resident population. The rescue plasmid is absent prior to the environmental change.

The intrinsic death rate of plasmid-free cells is 1 (in arbitrary units). Plasmids incur a metabolic cost *c*, which increases the death rate to 1 + *c*. Additionally, competition for resources between cells increases the net death rate of every cell type by a rate 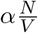, where *N* denotes the total population size. In the old benign environment, cells of the resident population divide at rate 1 + *s*_*p*_ with *s*_*p*_ *>* 0. After the environmental change, which occurs at time *t* = 0, the intrinsic division rate of cells of the resident population drops to 1 + *s*_0_ with *s*_0_ *<* 0 (e.g. through the application of a bacteriostatic antibiotic). This means that the resident population has a negative net growth rate even at low population densities and cannot persist in the new environment. ‘Rescue plasmids’ might enter the population through mutation of resident plasmids or through immigration from an external population as described further below. Cells carrying the rescue plasmid divide at rate 1 + *s*_*R*_ *>* 1. We assume that the intrinsic growth rate of these cells is positive (*s*_*R*_ − *c >* 0). They can thus rescue the population from extinction. The parameters are summarized in Table 1. In the main text, we assume that there is no segregational loss of plasmids, i.e. both daughter cells of a plasmid-carrying cell contain the plasmid. We discuss the effect of segregational loss in the Supplementary Information (see section S4 and Figures S6 and S7). In the Supplementary Information, we furthermore consider alternative models in which the cell type affects the intrinsic death rate rather than the intrinsic division rate and competition for resources and the metabolic cost of plasmid carriage reduce the cell division rate instead of the death rate (see section S1 and Figure S1). These modeling choices, which are partially arbitrary and partially for convenience, do not affect the results.

**Table 1:** Summary of the per-capita division and death rates of all cell types before and after the environmental change. Throughout the manuscript, we set *V* = 1.

|  | plasmid-free cells | resident plasmid cells | rescue plasmid cells |
| --- | --- | --- | --- |
| **before the environmental change** |  |  |  |
| death rate: | $1 + \alpha \frac{N}{V}$ | $1 + \alpha \frac{N}{V} + c$ | - |
| division rate: | $1 + s_p, s_p > c$ | $1 + s_p, s_p > c$ | - |
| **after the environmental change** |  |  |  |
| death rate: | $1 + \alpha \frac{N}{V}$ | $1 + \alpha \frac{N}{V} + c$ | $1 + \alpha \frac{N}{V} + c$ |
| division rate: | $1 + s_0, s_0 < 0$ | $1 + s_0, s_0 < 0$ | $1 + s_R, s_R > c$ |

We assume that both resident and rescue plasmids are conjugative. Horizontal transfer follows a mass-action model with transfer coefficients *β*_*r*_ and *β*_*R*_ respectively. E.g. resident plasmids transfer into plasmid-free cells at total rate 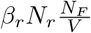. Resident and rescue plasmids can either be compatible or incompatible. If they are compatible, plasmids can transfer to cells that already contain a plasmid of the other type. Transfer of rescue plasmids then occurs at total rate 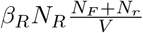. For simplicity, we do not treat cells with both plasmid types as a separate class; instead, we treat them as rescue plasmid cells and ignore the resident plasmid. This implies that (i) they do not carry an additional cost (ii) they can only act as donors of the rescue plasmid and (iii) that we can ignore transfer of resident plasmids into rescue plasmid cells. If the two plasmids are incompatible, we assume that they encode an entry or surface exclusion system, and transfer can thus only happen into plasmid-free cells, i.e. at total rate 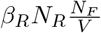 for the rescue plasmid. In the following, we set *V* = 1 (in arbitrary units) for simplicity and without loss of generality.

### 2.2 Outline of the analysis

We are interested in the probability that the population survives the environmental change and successfully adapts to the new conditions. Population survival is contingent on two factors – cells carrying the adaptive rescue plasmid need to appear in the population and they need to establish a long-term lineage of progenitor cells. If we know the rate of appearance of rescue plasmid cells *λ*(*t*) and the probability of establishment, i.e. non-extinction, *p*_est_(*t*) of a lineage founded by a rescue plasmid cell appearing at time *t*, we can obtain the probability of evolutionary rescue of a population *p*_res_:

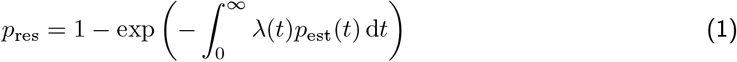

[see e.g. Orr and Unckless, 2008, Uecker et al., 2014, Tazzyman and Bonhoeffer, 2014].

In the following sections, we derive the rate of appearance of rescue plasmid cells *λ*(*t*) and their establishment probability *p*_est_(*t*).

### 2.3 Dynamics of the resident population

In the phase of interest, in which the population is at risk of extinction but has a chance for rescue, the rescue plasmid is rare (otherwise, the population would be safe) and the resident population is large (otherwise, the chance for rescue would be tiny). In our mathematical analysis, we can therefore study the dynamics of the resident population separately with a deterministic approach. Let us define the fraction of resident plasmid cells *φ*(*t*) = *N*_*r*_(*t*)*/N* (*t*). We consider three alternative models of increasing complexity for the dynamics of plasmid-free and plasmid carrying cells.

#### Model I - Constant *φ*

In the simplest model adapted from Tazzyman and Bonhoeffer [2014], we do not explicitly consider the dynamics of the resident plasmid, but rather we assume that the fraction of resident plasmid cells is constant in time, i.e. *φ*(*t*) = *φ* ∀*t*. In this model, *φ* is an exogenous parameter. The dynamics of the resident population simplify to a single differential equation

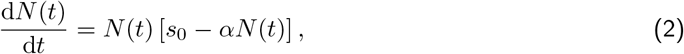

from which one can derive the dynamics of both cell types

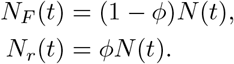

The initial number of cells is given by *N* (0) = *N*_0_, which is again an exogenous parameter.

#### Model II - Variable *φ*(*t*)

In the second model, we explicitly track the dynamics of the resident plasmid. The fraction of resident plasmid cells *φ*(*t*) is now an endogenous variable and is determined by horizontal transfers of the resident plasmid and cell division and death. The dynamics of the resident population are described by

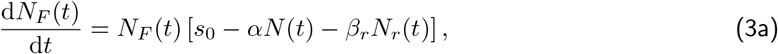

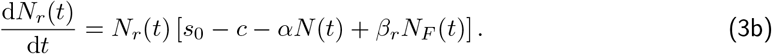

In Appendix A, we show that for non-trivial parameter values, *φ*(*t*) = *N*_*r*_(*t*)*/N* (*t*) is never constant, contrary to the assumption made in Model I. The models are only identical (i.e. *φ*(*t*) is constant) if *φ*(0) = 0 (absence of a resident plasmid), or *φ*(0) = 1 (all cells carry the resident plasmid), or *c* = 0 and *β*_*r*_ = 0 (non-transmissible resident plasmid with zero cost). In particular, for *c* = 0 (no plasmid cost) and *β*_*r*_ *>* 0, *φ*(*t*) always increases due to horizontal transfer. For *c >* 0, lim_*t→∞*_ *φ*(*t*) = 0; intuitively, at some point, the negative effect of plasmid cost becomes larger than the positive effect of horizontal transfer, as the population size and thus the opportunities for transfer decrease. However, it is possible that *φ*(*t*) temporarily increases after the environmental change before ultimately going to zero.

In this model, the initial fraction *φ*(0) = *φ*_0_ at the time of environmental change is still an exogenous constant parameter, namely, we do not make any assumption about the system before *t* = 0. The initial number of cells is again *N* (0) = *N*_0_.

In the main text, when comparing Models I and II, we set the constant fraction *φ* of resident plasmids in Model I equal to the initial fraction *φ*_0_ in Model II. In SI section S2, we make an additional model comparison, setting *φ* in Model I to the average fraction of *φ*(*t*) in Model II.

#### Model III - Variable *φ*(*t*) with pre-rescue dynamics

In the third model, we explicitly derive the initial population size *N* (0) and the initial fraction of resident plasmid cells *φ*(0) from the dynamics that occur before the environment changes at *t* = 0, the ‘pre-rescue dynamics’. Let *N*_*F,p*_ and *N*_*r,p*_ be the number of plasmid-free cells and resident plasmid cells during the pre-rescue dynamics. Then:

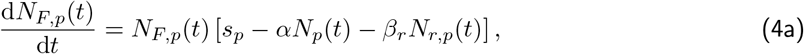

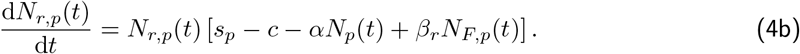

For the sake of simplicity, we assume that the system is in equilibrium at time *t* = 0, i.e. we use the equilibrium 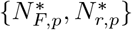 of Eq. (4) as the initial conditions of the dynamics determined by Eq. (3), 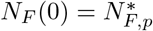 and 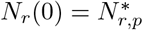 (illustrated in Figure B.1b).

We discard the trivial equilibrium 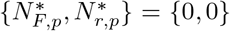. If 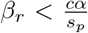, the only stable equilibrium is 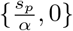, the ‘plasmid-free equilibrium’. If 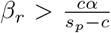, the only stable equilibrium is 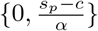, the ‘resident plasmid equilibrium’. Otherwise, for intermediate values of *β*_*r*_, only the ‘mixed equilibrium’ is stable 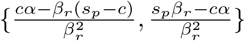. For the full analysis of the equilibria, see Appendix B.

Given these initial conditions, Model III is thus a special case of Model II. The reasoning presented in Model II about the dynamics of the fraction of resident plasmid cells *φ*(*t*) also applies in Model III after the environment changes. Namely, *φ*(*t*) will always shift away from its initial value 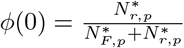 unless *φ*(0) = 0 or *φ*(0) = 1 (see Appendix A). Moreover, if the pre-rescue dynamics stabilizes to the ‘mixed equilibrium’, we show that the fraction of resident plasmid cells *φ*(*t*) always decreases after the environment changes (see Appendix B).

### 2.4 Origin of rescue plasmids

Following Tazzyman and Bonhoeffer [2014], we consider three ways in which the rescue plasmid may appear in the population. In Scenario A, the resident plasmid mutates to give rise to the rescue plasmid. An example of such evolution is mutation in *β*-lactamase genes that lead to resistance to a new *β*-lactam antibiotic, to which the original unmutated variant is sensitive [Salverda et al., 2010, San Millan et al., 2016]. In Scenarios B and C, the rescue plasmid enters the focal population via transfer from an external population. Transfer of an adaptive plasmid could, for example, happen from commensal to pathogenic bacteria during treatment [Schwalbe et al., 1990, Archambaud et al., 1991, Carattoli, 2003, Boekhoud et al., 2020] or between bacterial populations in polluted environments.

#### Scenario A - Rescue by de novo mutations on the resident plasmid

In the first model, the rescue plasmid appears by mutation from the resident plasmid. At replication of cells from the resident plasmid population (which happen at per-capita rate 1 + *s*_0_), the mutation occurs with probability *u*. With this, we obtain for the rate of appearance of the rescue plasmid

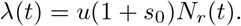

The resulting rescue plasmid is incompatible with the resident plasmid [Novick, 1987] and can thus only transfer into plasmid-free cells.

#### Scenario B - Rescue by immigration of an incompatible plasmid from an external population

In the second model, rescue plasmids appear in the focal population via horizontal transfer from an external population. The rescue plasmid is assumed to be incompatible with the resident plasmid and can hence only be transferred to plasmid-free cells:

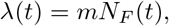

where *m* is the *per-capita* rate at which the rescue plasmid enters plasmid-free cells. Such a scenario could for instance happen if an external population has acquired the beneficial mutation or the entire beneficial gene in a similar environment, e.g. in the presence of the same antibiotic. If plasmid transfer between the source and the focal populations follows mass-action kinetics and the size of the source population *N*_source_ is constant, we can write *m* = *β*_source_*N*_source_, where *β*_source_ is the transfer coefficient from cells of the source population into cells of the focal population. Note that *N*_source_ is the number of cells that mix with the focal population and can be quite small, e.g. if there is low immigration of cells into the focal population from elsewhere.

In the main text, we use *m* as an independent parameter. This means that our results need to be interpreted as ‘rescue given a certain immigration rate (irrespective of what shapes *m*)’ or that the transfer rate *β*_source_ is entirely determined by the donor population and thus not correlated with the transfer coefficient *β*_*R*_ within the focal population. In Appendix C, we relax this assumption, considering *m* ∝ *β*_*R*_ (i.e. *β*_source_ ∝ *β*_*R*_).

#### Scenario C - Rescue by immigration of a compatible plasmid from an external population

Last, we assume that horizontal transfer from an external population can provide rescue plasmids that are compatible with the resident plasmid at rate *m*. Compatible plasmids can be transferred to both plasmid-free cells and resident plasmid cells. As pointed out above, for the sake of simplicity, we do not consider cells with both types of plasmids. I.e., when a rescue plasmid is transferred into a resident plasmid cell, the second becomes a simple rescue plasmid cell. For the rate of appearance, we have

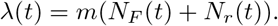

### 2.5 The establishment probability of the rescue plasmid

The rescue plasmid is initially rare, and stochasticity in the dynamics of the rescue plasmid cells can therefore not be ignored. The dynamics of a lineage founded by a single rescue plasmid cell can be modelled by a time-inhomogeneous birth and death process. Note that ‘birth’ here includes cell division as well as horizontal transfer since both are mathematically equivalent events. Thus, the birth rate *b* differs, depending on whether we consider a rescue plasmid that is incompatible (*b*^(*i*)^) or compatible (*b*^(*c*)^) with the resident plasmid. Since the population of rescue plasmid cells is small in the establishment phase, we can disregard competition between them in the death rate. The process thus becomes a branching process. Overall, we have:

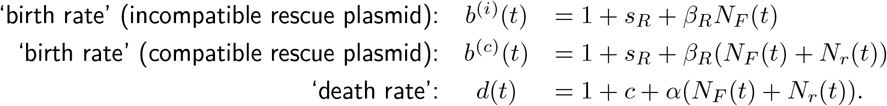

The establishment probability of a time-inhomogeneous birth and death process is known and given by

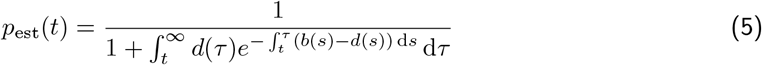

[Kendall, 1948, *Allen, 2010, Uecker and Hermisson, 2011, Tazzyman and Bonhoeffer, 2014]*. *(Note that here and elsewhere ‘dt*’ with an upright ‘d’ is the differential, not to be mixed up with the death rate function *d*(*t*).)

For constant birth and death rates (i.e. for a population of constant size and *φ* = const.), the establishment probability of an adaptive plasmid that is incompatible with the resident plasmid simplifies to

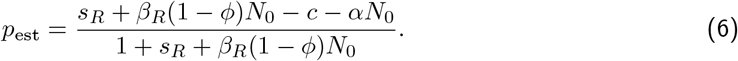

### 2.6 Numerical analysis and simulations

We could not derive closed-form solutions for the rescue and establishment probabilities, Eq. (1) and (5). We provide numerical results for which the upper limits of integration in (1) and (5) are not infinite but large values, respectively *T*_res_ and *T*_est_:

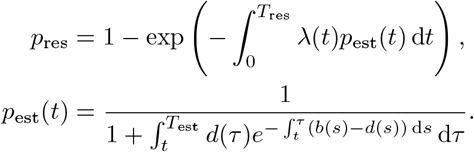

We choose *T*_res_ such that in Model I, the resident population size is reduced to a single cell, i.e. *N* (*T*_res_) = 1. From equation (2), we can show that

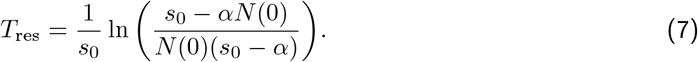

In Models II and III, the resident population size decreases faster than in Model I due to the plasmid cost *c* such that with *T*_res_ given in equation (7), we have *N* (*T*_res_) ≤1 in these models.

We choose *T*_est_ = 2*T*_res_ so that we still track the dynamics of rescue plasmid cells appearing just before *T*_res_ for a time of *T*_res_ and do not consider them established prematurely. Larger values for the limits of integration *T*_res_ and *T*_est_ do not visibly change the results.

In addition to the numerical analysis, we also run stochastic simulations of the full stochastic model using the Gillespie algorithm. Simulation results are the average result over 10^4^ simulations. When the pre-rescue dynamics are considered (Model III), we use the equilibrium values previously derived as the starting point for stochastic simulations in the rescue phase. Each simulation stops either (i) when the total population goes to extinction *N*_*F*_ + *N*_*r*_ + *N*_*R*_ = 0, or (ii) when the number of rescue plasmid cells reaches a threshold *N*_*M*,max_. We choose this threshold such that a population of *N*_*M*,max_ rescue plasmid cells does not go to extinction with probability 99% in the absence of the other types and competition. If rescue plasmid cells are the only type, their dynamics simplify to a time-homogeneous birth and death process with

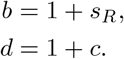

The establishment probability of a population of size *n* of such a branching process is 1 − (*d/b*)^*n*^ [Allen, 2010]. Thus, the threshold *N*_*M*,max_ is defined by

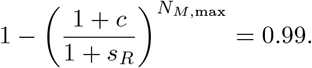

We can check that neglecting competition between rescue plasmid cells is justified: for all values of parameters *s*_*R*_ and *c* that we explore, the threshold *N*_*M*,max_ is lower than 100. The value for the competition parameter *α* that we use in the remaining of the paper is 10^−6^. Thus, the net competition rate *αN*_*M*,max_ is negligible when the stopping criterion is met, 1 + *sM ≫ αN*_*M*,max_ and 1 + *c ≫ αN*_*M*,max_. There could still be competition with other cell types. However, simulation results with higher thresholds in the stopping criterion were not visibly different.

Unless stated otherwise, we use the following parameters in the figures: *s*_0_ = −0.1; *s*_*R*_ = 0.1; *s*_*p*_ = 0.1; *c* = 0; *α/V* = 10^−6^; *N*_0_ = 10^6^; *u* = 10^−6^; *m* = 10^−6^. Similar to Tazzyman and Bonhoeffer [2014], we consider a large range of transfer coefficients, ranging from 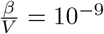 up to 10^−4^. We do not consider lower values of *β/V* than 10^−9^ since, as can be seen in the figures below, rescue probabilities converge for low *β/V* and conjugation becomes irrelevant for rescue at these low values. We put this range into the context of empirically measured transfer coefficients in the Discussion section. We remind the reader that we are setting *V* = 1 throughout he manuscript.

### 2.7 Supplementary remarks

Before presenting the results, we would like to state a few general observations that directly follow from Eq. (1) and the methods used for the analysis. From Eq. (1), we see that the rescue probability increases with the initial population size *N*_0_, the mutation probability *u*, and the immigration rate *m*. Changes in any of those parameters do not affect how the rescue probability depends on the transfer coefficients. The choice of specific values does therefore not influence our results qualitatively.

Finally, since we model the dynamics of the resident population deterministically in the analysis, we can obtain the rescue probability of an *M* times larger bacterial population occupying an *M* times larger volume by

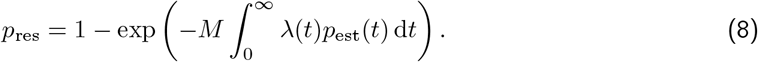

## 3 Results

In the main text, we assume for all models that the horizontal transfer coefficients of the resident and the rescue plasmid are identical, i.e. *β*_*r*_ = *β*_*R*_ = *β*. This is especially likely when the rescue plasmid is a mutated variant of the resident plasmid, either being generated through *de novo* mutation *in situ* or entering the population from the outside. In SI section S3, we study two alternative assumptions: transfer coefficients that are proportional to each other, reflecting a host contribution to the transfer coefficient, and independent transfer coefficients.

As shown by Eq. (1), the rescue probability is determined by the interplay between the appearance of rescue plasmid cells *λ*(*t*) and their establishment probability *p*_est_(*t*). Competition for host cells between the resident and the rescue plasmid affects both quantities. Understanding how *λ*(*t*) and *p*_est_(*t*) are influenced by plasmid competition is essential for understanding the dependency of the rescue probability on the transfer coefficient *β*. Before we consider rescue probabilities, we therefore illustrate this influence in the simplest model version (Model I), where the fraction of resident plasmids *φ* is constant (Figure 2), which is essentially a recapitulation of the arguments made by Tazzyman and Bonhoeffer [2014].

**Figure 2.**
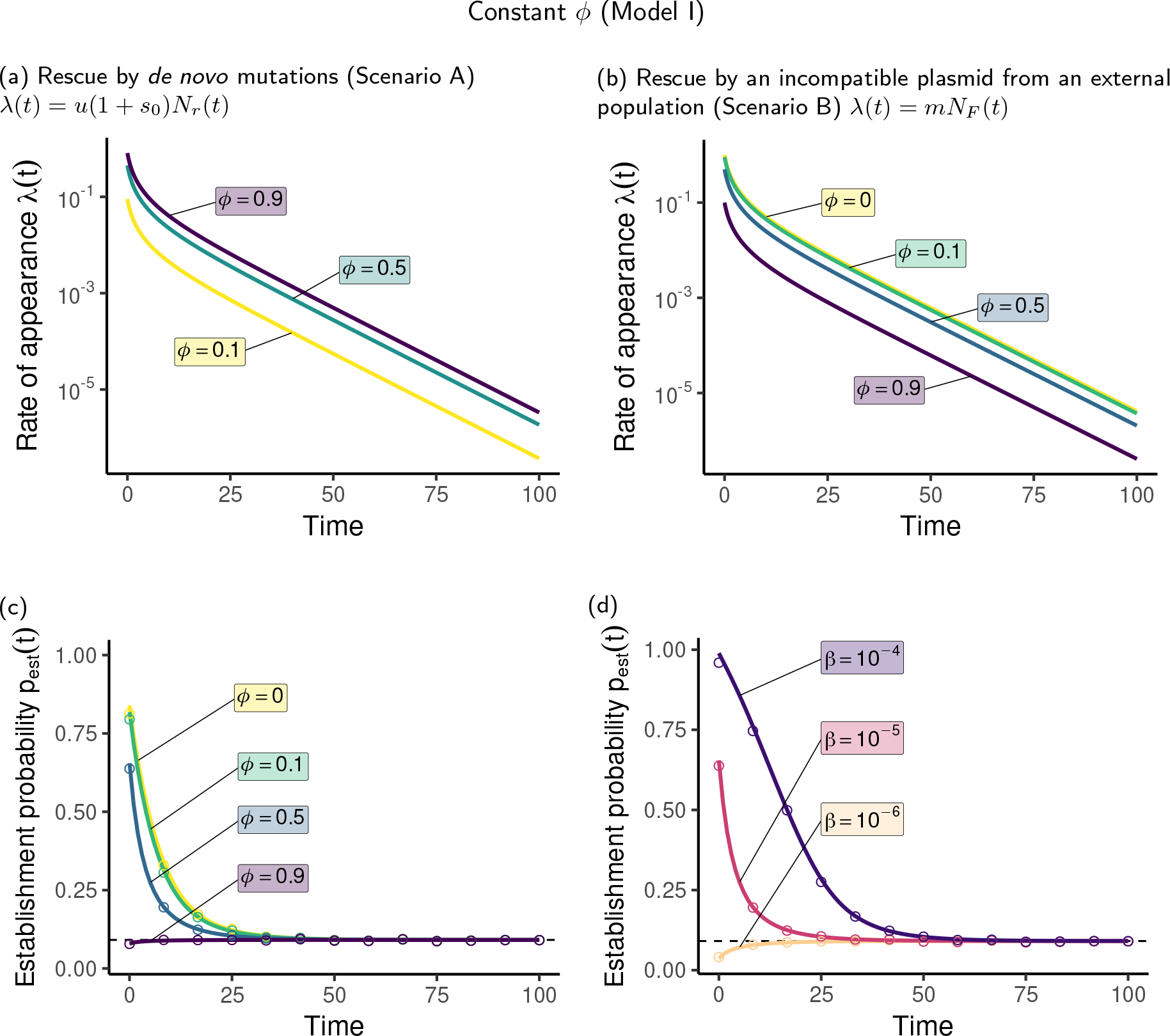
Rate of appearance and establishment probability of the rescue plasmid in Model I, in which the fraction of resident plasmid cells *φ* is an exogenous parameter. Panels **(a)** and **(b)** show the rate of appearance for multiple values of the fraction *φ*. In Panel (a), the rescue plasmid is generated through *de novo* mutations on the resident plasmid. The rate of appearance decreases over time since the sub-population of resident plasmid cells decreases. The higher the fraction *φ* of resident plasmid cells, the higher the mutational input. In Panel (b), the rescue plasmid enters the focal population through immigration and is incompatible with the resident plasmid. The rate of appearance decreases over time here as well since the sub-population of plasmid-free cells that can receive the rescue plasmid decreases. However, in contrast to the scenario in Panel (a), the rate of appearance is higher for lower fractions of resident-plasmid cells. Panels **(c)** and **(d)** show the establishment probability of a rescue plasmid appearing at time *t* for multiple values of the fraction of resident plasmid cells *φ* and the horizontal transfer coefficient *β*. The dashed line indicates the establishment probability in the absence of resident cells: 1 − (1 + *c*)*/*(1 + *s*_*R*_). The establishment probability either decreases over time (as a result of a decrease in the absolute number of plasmid-free cells that are targets for horizontal transfer, as the the total population size decreases over time) or slightly increases over time (as a result of relaxed competition). The establishment probability is high if resident plasmid cells are rare and the transfer coefficient high. Circles indicate stochastic simulation results. Parameters (if not indicated otherwise in the figure): *β* = 10^−5^, *φ* = 1*/*2.

If the rescue plasmid appears through *de novo* mutations, the rate of appearance increases with *φ* since resident plasmid cells are needed to generate mutants (Figure 2a). In contrast, if the rescue plasmid enters the population from the outside and is incompatible with the resident plasmid, the rate of appearance decreases with *φ* (Figure 2b). For a compatible plasmid coming from an external population, finally, *λ*(*t*) is independent of the fraction of resident plasmids in the population (not shown). For incompatible rescue plasmids, the presence of resident plasmid cells inhibits their establishment probability because it decreases the pool of plasmid-free cells available for horizontal transfer (Figure 2c). The interference of the resident plasmid with establishment of the rescue plasmid can also be seen in the simplified equation (6). The effect becomes less important over time since the population of plasmid-free cells goes to extinction and the contribution of horizontal transfer dwindles (Figure 2c). The transfer coefficient of rescue plasmids *β* favours establishment but its effect also decreases in time for the same reason (Figure 2d). Note that, when horizontal transfers of rescue plasmids are rare, the establishment probability increases through time due to the reduction in competition for resources as the resident population goes to extinction. This can be the case when not enough plasmid-free cells are available (*φ* = 0.9 in Figure 2c) or when transfer coefficients are too low (*β* = 10^−6^ in Figure 2d).

In the next three sections, we first consider rescue by *de novo* mutations, then by immigration of an incompatible plasmid, and finally by immigration of a compatible plasmid.

### 3.1 Scenario A - Rescue by de novo mutations on the resident plasmid

In the simple model with constant *φ* (Model I), a higher transfer coefficient of rescue plasmids *β* always increases the rescue probability (Figure 3a). As outlined above, the fraction of resident plasmid cells *φ* has two opposing effects on the rescue probability – having more resident plasmids in the population is on the one hand beneficial since it increases the rate of appearance; on the other hand, it hampers the spread of incompatible rescue plasmids. As explained by Tazzyman and Bonhoeffer [2014], the net effect depends on the transmission coefficient of the rescue plasmid. If transfer is unlikely (low *β*), the beneficial effect on *λ*(*t*) always outweighs the negative effect on *p*_est_(*t*). However, for intermediate and high values of *β*, the effect of *φ* is non-linear, i.e. there is an intermediate optimal value of *φ* which maximizes the compromise between maintaining a large rate of appearance and preventing competition at the plasmid-level (Figure 3a). As *β* increases, the maximum shifts to ever higher values of *φ* (see Figure 1 in Tazzyman and Bonhoeffer [2014]).

**Figure 3.**
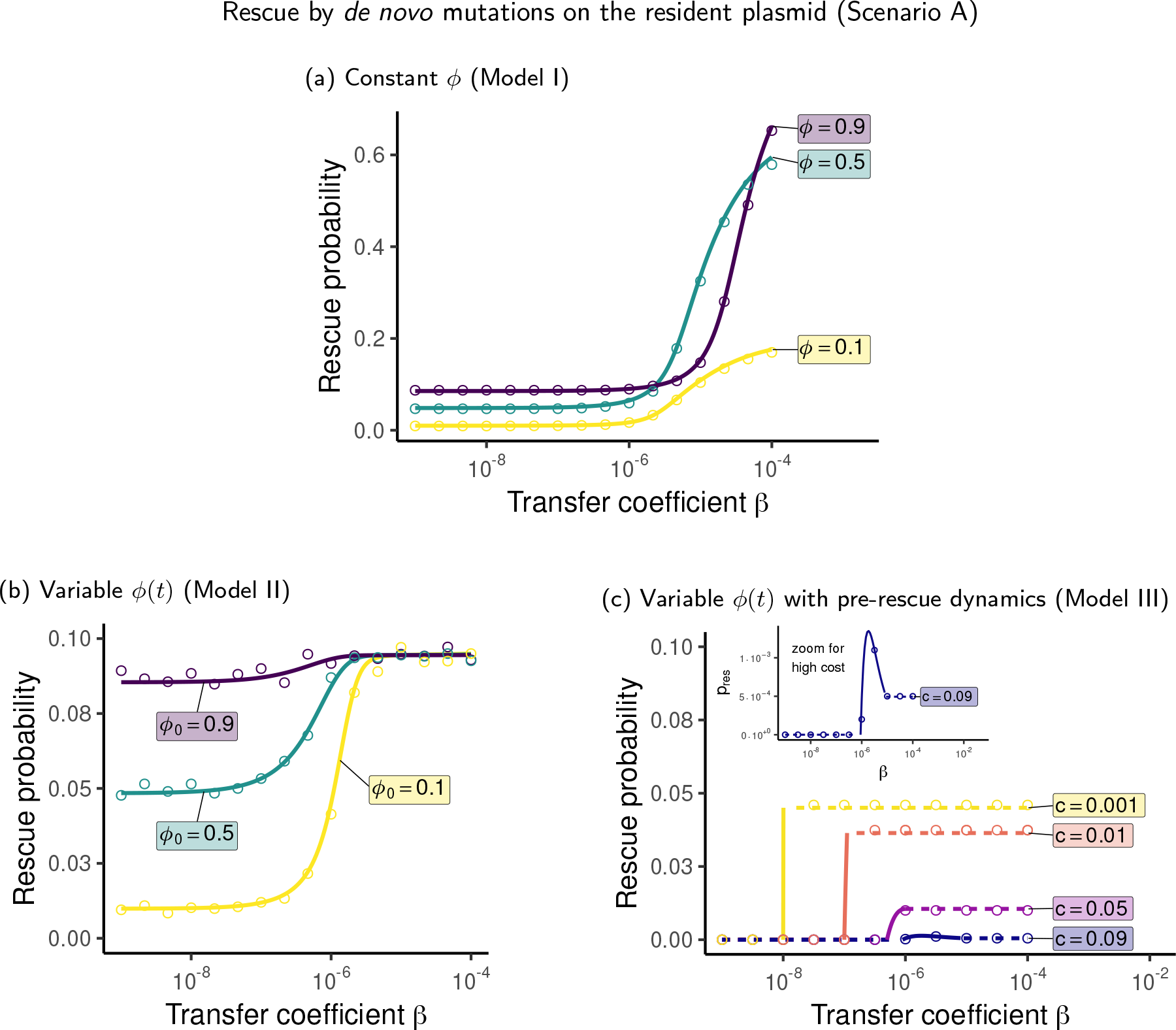
Probability of evolutionary rescue by *de novo* mutations as a function of the transfer coefficient for all three model variants. In Panel **(a)**, the fraction *φ* of resident plasmid cells is an exogenous parameter and constant in time. In Panel **(b)**, the dynamics of resident plasmid cells are explicitly modeled, and their fraction *φ*(*t*) in the population changes over time. The initial fraction *φ*(0) = *φ*_0_ is an exogenous parameter. In Panel **(c)**, the dynamics of the resident plasmid cells are explicitly modeled both before and after the environmental change. In panel (c), solid lines indicate the range of *β* for which the population consists of a mix of plasmid-free and resident plasmid cells prior to the environmental change, while dashed lines indicate ranges of *β* for which the population consists of only plasmid free cells (low *β*) or only resident plasmid cells (high *β*). In all panels, the probability of rescue increases with the transfer coefficient with the exception of plasmids with high cost *c* in panel (c).

Figure 3b shows the rescue probability when we relax the assumption that the fraction of resident plasmid cells *φ*(*t*) is constant (Model II). In this case, the parameter *β* acts on the transfer rate of both resident and rescue plasmids. Larger values of *β* lead to a higher number of resident plasmid cells and thus increase the rate of appearance. The net rate at which a focal rescue plasmid cell transfers its plasmid to plasmid-free cells is *βN*_*F*_. Therefore, larger values of *β* (i) directly increase the net rate of transfer and (ii) indirectly decreases the net rate via competition at the plasmid-level, since fewer plasmid-free cells are available. To be able to compare the results with Model I, we show results for *c* = 0 (no plasmid costs). Overall, a higher transfer coefficient *β* seems to be beneficial for rescue; for large *β*, the rescue probability eventually converges to a constant value that is independent of *φ*_0_ (Figure 3b). For high plasmid costs, the dependency of the rescue probability on *β* turns non-monotonic, since large values of *β* drive the population to a state with few plasmid-free cells and many resident plasmid cells that die rapidly with little chance of producing mutants (see Figure S9a).

As discussed above (see also Appendix A), the fraction of resident plasmid cells *φ*(*t*) never stays constant when we allow it to change (Figure 4a). If we compare the predictions of Model I (with *φ* = *φ*_0_) and Model II (Figure 4b), we see that Model I strongly overestimates the rescue probability for high *β*, since it underestimates the extent of the competition at the plasmid-level. The predictions of Models I and II converge for low values of *β*. This is because for low *β* (and no cost, *c* = 0, as assumed here), the fraction of resident plasmid cells moves away only slowly from its initial value *φ*_0_ in Model II and horizontal transfer is of minor importance for establishment. Interestingly, for intermediate values of *β*, Model I underestimates the rescue probability, especially for small *φ*_0_. This most likely comes from the fact that in Model II only, higher values of *β* cause an increase in the rate of appearance. This favors rescue for intermediate values of *β* and low *φ*_0_ because the marginal benefit on rescue of increasing the rate of appearance is larger than the marginal cost of plasmid competition under these conditions.

**Figure 4.**
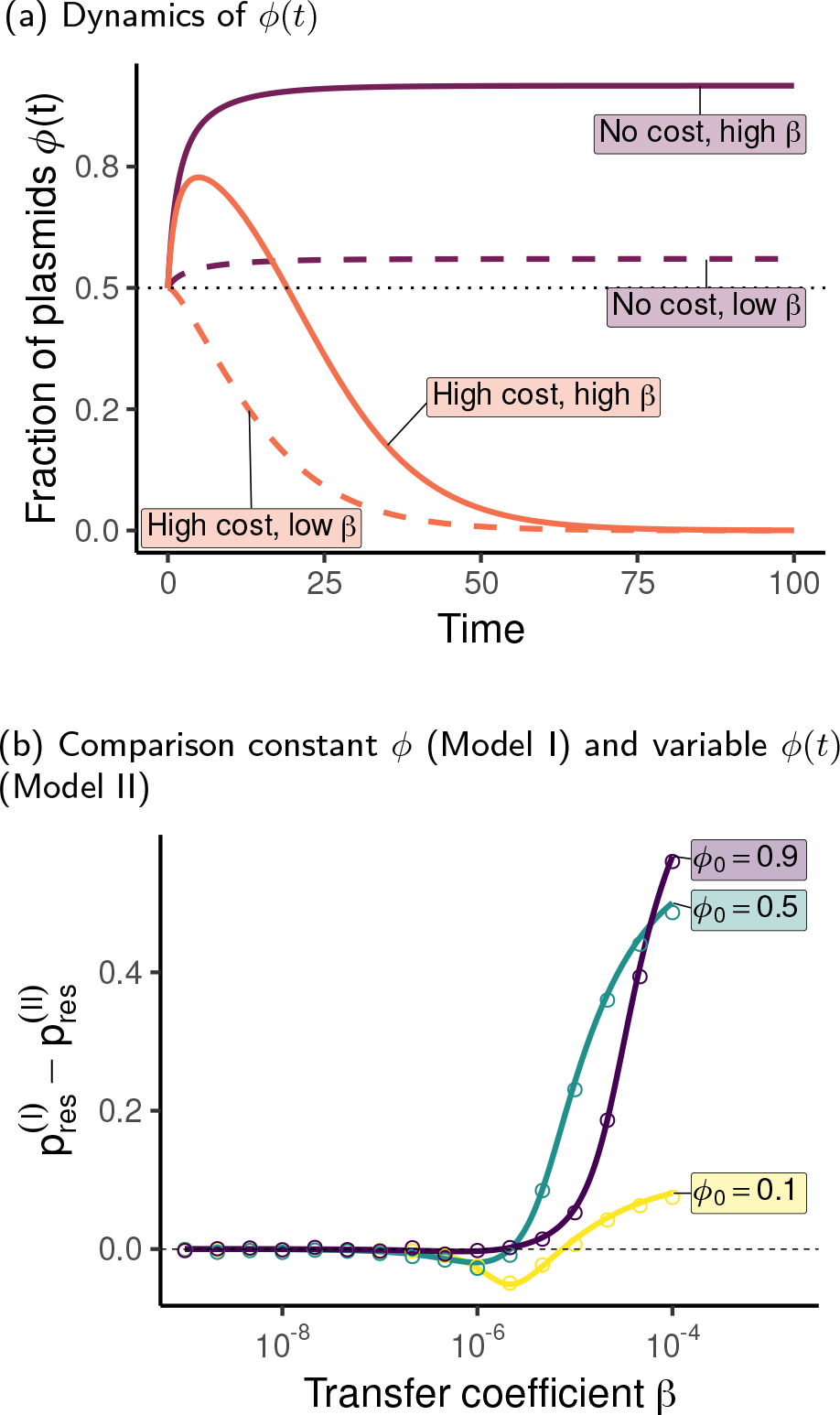
Comparison of the models with a constant fraction *φ* of resident plasmid cells (Model I) and explicitly modeled dynamics *φ*(*t*) with a given initial value *φ*_0_ (Model II). In Panel **(a)**, the solid and dashed lines show the dynamics of *φ*(*t*) starting with *φ*_0_ = 0.5 for no costs (*c* = 0) and high costs (*c* = 0.1) and for low (*β* = 10^−7^) and high transfer coefficients (*β* = 10^−6^). The dotted line corresponds to a constant of *φ* = 0.5. Panel **(b)** shows the difference in the probability of rescue by *de novo* mutations (Scenario A) between Model I with constant *φ* and Model II with variable *φ*(*t*) as a function of the transfer coefficient *β*. Positive values mean that the simpler Model I overestimates the rescue probability compared to the more realistic Model II.

Finally, we can determine the initial frequency of resident plasmid cells *φ*_0_ from the pre-rescue dynamics (Model III). This case is similar to Model II with the difference that *β* plays a role also in the pre-rescue dynamics, determining the prevalence of plasmid-free and resident plasmid cells at the equilibrium (represented in Figure B.1a). Three cases can be distinguished. If the population is in the ‘plasmid-free equilibrium’ at the time of environmental change, then the rate of appearance during the rescue phase is null *λ*(*t*) = *u*(1 + *s*_0_)*N*_*r*_(*t*) = 0, dashed lines in the range of low *β* in Figure 3c. If the population is in the ‘resident plasmid equilibrium’, no plasmid-free cells are present during the rescue phase, hence *β* has no impact, dashed lines in the range of high *β* in Figure 3c. For intermediate values of *β*, the population is in the ‘mixed equilibrium’ with plasmid-carrying and plasmid free cells present, solid lines in Figure 3c. Note that the extent of this intermediate range 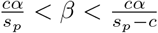 depends on the plasmid cost *c* (see Figure B.1a). We show in Appendix B that for low plasmid costs *c*, the number of resident plasmid cells at the time of environmental change *N*_*r*_(0) always increases with the transfer coefficient *β*. Yet, for high values of *c*, high values of *β* can actually decrease the initial number of resident plasmid cells *N*_*r*_(0). Intuitively, if a plasmid is too costly and virulent, it can harm its host and thus its own growth, such as parasites with high levels of virulence [Frank, 1996]. As a consequence, the transfer coefficient *β* can have a negative impact on the rescue probability if the plasmid cost is high enough (Figure 3c see inset).

### 3.2 Scenario B - Rescue by immigration of an incompatible plasmid from an external population

When incompatible rescue plasmids come from an external population via horizontal transfer (Scenario B), their rate of appearance is proportional to the number of plasmid-free cells *N*_*F*_ (*t*) (see section 2.4).

Figure 5 displays the rescue probability for Scenario B and for the three different assumptions regarding the dynamics of the resident population, Models I-III. If the fraction of resident plasmid cells *φ* is constant (Model I), higher values of *β* only imply higher rates of rescue plasmid transfers as before, leading to an increase in the rescue probability with *β* (Figure 5a). A large prevalence of resident plasmids *φ* is always detrimental, since they hinder both the appearance of rescue plasmid cells *λ*(*t*) and their chances of escaping stochastic loss *p*_est_(*t*).

**Figure 5.**
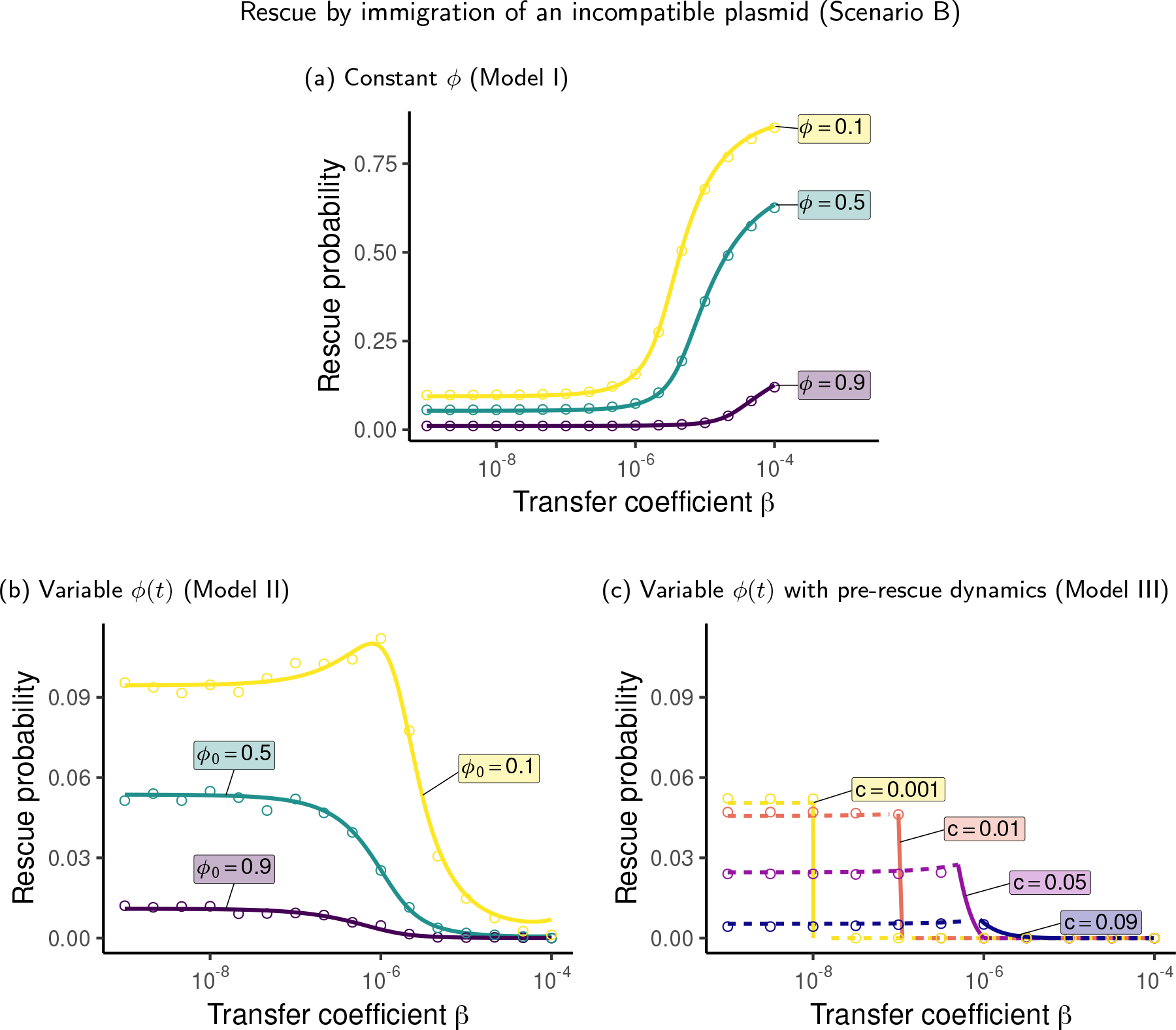
Probability of evolutionary rescue by immigration of an incompatible plasmid that has the same transfer coefficient as the resident plasmid as a function of the transfer coefficient for all three model variants. In Panel **(a)**, the fraction *φ* of resident plasmid cells is an exogenous parameter and constant in time. In Panel **(b)**, the dynamics of resident plasmid cells are explicitly modeled, and their fraction *φ*(*t*) in the population changes over time. The initial fraction *φ*(0) = *φ*_0_ is an exogenous parameter. In Panel **(c)**, the dynamics of the resident plasmid cells are explicitly modeled both before and after the environmental change. In panel (c), solid lines indicate the range of *β* for which the population consists of a mix of plasmid-free and resident plasmid cells prior to the environmental change, while dashed lines indicate ranges of *β* for which the population consists of only plasmid free cells (low *β*) or only resident plasmid cells (high *β*). In Models II and III, a higher transfer coefficient mostly reduces the chances of rescue.

When we consider that resident plasmids transfer with coefficient *β* within the focal population, leading to changing *φ*(*t*) (Model II), the fact that the rescue plasmid can only immigrate into plasmid-free cells drastically changes the predictions compared to the scenario of rescue by *de novo* mutations. The effect of increasing horizontal gene transfer on the rescue probability becomes almost systematically negative since it leads to higher numbers of resident plasmids cells (Figure 5b). Only when the initial fraction of resident plasmid cells *φ*_0_ and the transfer coefficient *β* are low, can an increase in *β* result in higher chances of rescue. Indeed, when plasmid free cells *N*_*F*_ (*t*) are very common, the marginal benefit of increasing the transfer rate of rescue plasmids *βN*_*F*_ (*t*) is large enough to outweigh the marginal costs of reducing the rate of appearance *λ*(*t*) = *mN*_*F*_ (*t*).

When pre-rescue dynamics are taken into account (Model III), the effect of *β* depends on the equilibrium at the time of change. If only plasmid-free cells are present at the onset of the rescue phase, the only effect of *β* is to increase the spread of the rescue plasmid, but this effect is weak since *β* is low in that regime, dashed lines in the range of low *β* in Figure 5c. If a mix of plasmid-free and resident plasmid cells are present, larger values of *β* increase the fraction of resident plasmid cells, diminishing the rate of appearance during the rescue phase *λ*(*t*) = *mN*_*F*_ (*t*) and increasing plasmid competition for hosts, which ultimately outweighs the beneficial direct effect of rescue plasmid transfer on establishment, solid lines in Figure 5c. Otherwise, if prior to change all cells carry the resident plasmid (‘resident plasmid equilibrium’), the rate of appearance and thus the rescue probability are null, dashed lines in the range of high *β* in Figure 5c.

So far, we made the assumption that the *per-capita* immigration rate of the rescue plasmid *m* is independent of *β*. In Appendix C, we consider the probability of rescue when the immigration rate is proportional to the transfer coefficient, *m* ∝ *β*, reflecting the contribution of the plasmid on its transfer. This provides an additional direct advantage of increasing *β*. As a consequence, unlike for *m* = const., the rescue probability increases with *β* over a considerable range, making the trends in Models I and II more similar (Fig. C.1a and c and Fig. C.2). Yet, for large *β*, the negative effect of high transfer rates still dominates, and the rescue probability starts to decrease with increasing *β* (Fig. C.1).

### 3.3 Scenario C - Rescue by immigration of a compatible plasmid from an external population

In Scenario C, rescue plasmids that are compatible with the resident plasmid immigrate from an external population. The rate of appearance and the rate at which rescue plasmids are transferred inside the focal population are proportional to the total number of cells in the resident population *N* (*t*) = *N*_*F*_ (*t*) + *N*_*r*_(*t*) (see section 2.4).

In Model I, unlike in the previous scenarios, the fraction of resident plasmid cells *φ* does not have any effect on the rescue probability as shown in Figure 6, since the exact composition of the resident population is irrelevant. A higher transfer coefficient *β* increases the establishment probability and therefore the rescue probability (Figure 6a).

**Figure 6.**
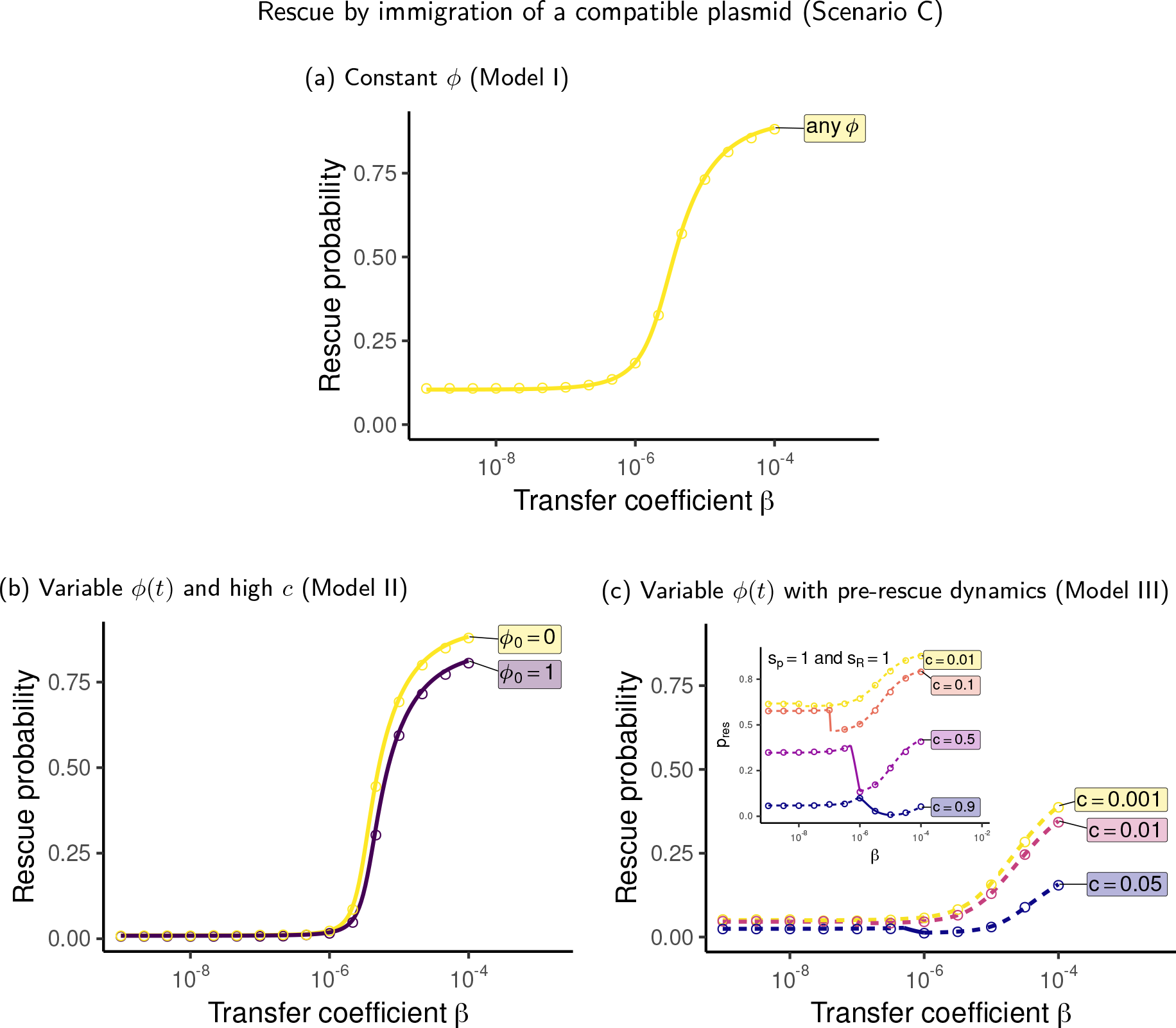
Probability of evolutionary rescue by immigration of a compatible plasmid that has the same transfer coefficient as the resident plasmid as a function of the transfer coefficient for all three model variants. In Panel **(a)**, the fraction *φ* of resident plasmid cells is an exogenous parameter and constant in time. In that case, the fraction of resident plasmid cells does not affect rescue. In Panel **(b)**, the dynamics of resident plasmid cells are explicitly modeled, and their fraction *φ*(*t*) in the population changes over time. The initial fraction *φ*(0) = *φ*_0_ is an exogenous parameter and the plasmid carries a cost *c* = 0.09. When the plasmid is costly (*c >* 0), the total resident population decreases faster if the fraction of resident plasmid cells is higher. This leads to a slightly lower rescue probability for *φ*_0_ = 1 than for *φ*_0_ = 0. In Panel **(c)**, the dynamics of the resident plasmid cells are explicitly modeled both before and after the environmental change. In panel (c), solid lines indicate the range of *β* for which the population consists of a mix of plasmid-free and resident plasmid cells prior to the environmental change, while dashed lines indicate ranges of *β* for which the population consists of only plasmid free cells (low *β*) or only resident plasmid cells (high *β*). If the plasmid imposes a cost on the bacterial host and pre-rescue dynamics are taken into account, the dependency of the rescue probability on the transfer coefficient can become non-monotonic. This effect is better seen in the inset graph with higher values for the fitness parameters *s*_*p*_ = *s*_*R*_ = 1.

Since only the total resident population size *N* (*t*) matters and not its composition *φ*, the same result holds when one relaxes the assumption that *φ*(*t*) is constant (Model II), as long as the plasmid cost is null (result not shown, same as in Figure 6a). If plasmids impose a fitness cost *c >* 0, the rescue probability still increases with *β* but decreases with the initial fraction *φ*_0_ of resident plasmid cells. With *c >* 0, the higher the prevalence of resident plasmids in a population the faster its decline and, thus, the lower the rate of appearance of rescue plasmids. The effect on the establishment probability is more subtle since the total resident population size drives both the competition for resources *αN* (*t*) and the rate at which a rescue plasmid cell transfers its plasmid *βN* (*t*). Figure 6b illustrates the interplay between these effect for two values of the initial fraction of resident plasmid cells *φ*_0_.

Figure 6c displays the effect of *β* when pre-rescue dynamics are considered. If the population is either in the ‘plasmid-free equilibrium’ or in the ‘resident plasmid equilibrium’ at the time of environmental change, then the fraction of resident plasmid cells will be constant during the rescue phase, respectively *φ*(*t*) = 0 and *φ*(*t*) = 1. The transfer coefficient *β* only acts on the establishment probability of rescue plasmid cells, hence increasing the likelihood of rescue, dashed lines in Figure 6c. On the contrary, when the population is in the ‘mixed equilibrium’, larger values of *β* lead to a larger fraction of resident plasmids cells *φ*(0) when the environment changes (Eq. (B.3)). In this case, it turns out that the negative effect on the rate of appearance (due to the decreased population size with a costly plasmid; Eq. (B.2)) is larger than the positive effect on the establishment probability, solid lines in Figure 6c.

Results for *m* ∝ *β* can be found in Appendix C.

## 4 Discussion

In this article, we proposed a set of models derived from Tazzyman and Bonhoeffer [2014] to systematically study the limits to evolutionary rescue on a conjugative plasmid in the presence of a resident plasmid. The rescue plasmid can arise via *de novo* mutation on the resident plasmid (Scenario A) or immigrate from an external population, in which case in can be either incompatible or compatible with the resident plasmid (Scenarios B and C, respectively). Crucially, for the main part of the manuscript, we assume that the two plasmid types transfer at the same rates, which is especially relevant if the rescue plasmid is a variant of the resident plasmid. We step-by-step made the model more complex: in Model I, the fraction of resident plasmids is assumed to remain constant over time; in Model II, we explicitly consider the dynamics of the resident plasmid after the environmental change; in Model III, finally, we also study the dynamics prior to change. Horizontal transfer becomes only relevant for rescue when the transfer coefficient is sufficiently large (Fig. 3, 5, 6); low transfer coefficients do not visibly affect the rescue probability (unless the immigration of the rescue plasmid increases with the transfer coefficient, see Fig. C.1). In the presence of plasmid competition and plasmid costs, increasing the rate of horizontal gene transfer has various, sometimes opposing effects on plasmid-driven rescue. Depending on which of them dominates, the net effect can be decreasing, increasing, or non-monotonic.

When the fraction of resident plasmid cells *φ* is constant in time (Model I), our results coincide with those by Tazzyman and Bonhoeffer [2014], which is expected since the models are identical except for the inclusion of resource competition in our model. Higher transfer coefficients *β* always promote the establishment of rescue plasmid cells, and thus promote rescue. However, in reality, the fraction of resident plasmid cells *φ* is determined by the interplay of conjugation, plasmid costs, and segregational loss. We showed that during population decline, the fraction *φ*(*t*) always moves away from its initial value *φ*(0) (see Figure 4a and Appendix A). If we take this into account (Model II), the role of horizontal gene transfer on rescue becomes more complex. A higher transfer coefficient leads to a larger number of resident plasmid cells. Besides the direct positive effect on the spread and thus the establishment probability of the rescue plasmid, the transfer coefficient thus also plays a similar role as the (fixed) fraction of resident plasmid cells in Model I. We find that in the interplay of the various effects, higher rates of transfer benefit rescue if rescue is based on new mutations on the resident plasmid unless plasmid costs are high, in which case the dependency becomes non-monotonic. At least in the absence of plasmid costs, the qualitative predictions of Model II thus agree with those in Model I but rescue probabilities are often much lower. In contrast, in Scenario B – rescue by immigration of an incompatible plasmid – predictions strongly diverge, and Model II predicts mostly a decreasing trend in the rescue probability with the transfer coefficient. If the two plasmids are compatible, the rescue probability increases with *β* in Model II even if plasmid costs are high.

We provide a further comparison between Models I and II in SI section S2, where we set the fraction of resident plasmid cells in Model I to its average in Model II (rather than to *φ*_0_ as in the main text). This means that *φ* remains constant over time in Model I but is dependent on *β*. With this parameterization in Model I, predictions of the two models become similar, both qualitatively and quantitatively. The discrepancies between the models that we observe in the main text are thus mainly driven by the change in the average value of the fraction of resident plasmid cells rather than the specific time dynamics. One should also note that for all scenarios, Model I and II converge for low transfer coefficients *β*, where the effect of transfer is generally weak.

The results of Model II and III are mostly similar regarding the overall trends. But we find, for example, that for Model III and strong plasmid costs, there is a range of *β* in which the rescue probability drops with the transmission coefficient due to the negative effect on the population size (Fig. 3c, Fig. 6c). In the Supplementary Information, we show that our results are robust to changes in the way the fitness parameter *s*, the plasmid cost *c*, and the strength of competition for resources *α* affect the birth and death rates of rescue plasmid cells (SI section S1). We also investigate the effect of plasmid segregation which could potentially change the dynamics since rescue and resident plasmid cells produce new plasmid-free cells at a constant rate. We show that high rates of plasmid segregation usually reduce the probability of rescue but do not qualitatively change our results (SI section S4). Interestingly, in some specific circumstances, plasmid segregation can make rescue more likely when plasmid-free cells are needed to host external incompatible plasmids (Scenario B).

Note that in the main text, we made the assumption that the rate *m* at which rescue plasmids are being transferred from an external population (Scenarios B and C) is independent of the rate *β* at which they are then being transferred within the focal population. This means that the transfer rate *m* is fully determined by properties of the external population. In Appendix C, we present an extension of our model where the immigration rate *m* is influenced by plasmid-related properties. In that case, the rate *m* and the transfer coefficient *β* are correlated. We show that this additional positive effect of higher *β* overall increases the circumstances under which an increase in *β* benefits rescue, but cannot always compensate for the negative impact of higher *β* on rescue via the reduction of plasmid-free cells (Figure C.1).

Since the transfer coefficient *β* is the central parameter of our study, it is worth pausing here to briefly discuss how often plasmids conjugate. The rate of conjugation depends on a multitude of factors including the host and the donor strain, the plasmid – especially whether conjugation is repressed or derepressed –, the environment, and the presence and absence of co-segregating plasmids [Dionisio et al., 2002, Gama et al., 2018, Alderliesten et al., 2020, Sheppard et al., 2020]. Measured conjugation efficiencies vary over many orders of magnitudes [Dionisio et al., 2002, Alderliesten et al., 2020, Sheppard et al., 2020]. Surprisingly, no consensus method for measurement exists though [Huisman et al., 2022, Kosterlitz et al., 2022, Kosterlitz and Huisman, 2023]. Many studies do not estimate *β* but the number of transconjugants per donor or per recipient after a given time. A recent meta-analysis by Sheppard et al. [2020] found that transfer coefficients (our parameter *β*) lie between 1.6·10^−20^ and 4.8·10^−7^ ml cells^-1^ h^−1^. Conjugation can be more frequent *in vivo* than *in vitro* [Göttig et al., 2015, Ding et al., 2022]. Measuring conjugation in the chicken gut, Fischer et al. [2019] estimated a conjugation coefficient of 10^−4^ cells^-1^ h^−1^ (note the absence of volume here) but with high uncertainty and caveats. Bacterial blooms during inflammation were found to foster high rates of transfer in the gut [Stecher et al., 2012, 2013]. Throughout the manuscript, we have given our parameters in arbitrary units. To set them in the context of measurements, we need to specify units. In the following, we provide a few examples. For *V* = 1 mL and measuring time in hours (i.e. the natural death rate of cells would be 1 h^−1^), our considered range of *β* would be 10^−9^ - 10^−4^ ml cells^-1^ h^−1^ and thus exceed the range that is usually measured by three orders of magnitude. For smaller volumes *V*, our range corresponds to the higher end of measured values (e.g., *β* between 10^−11^ and 10^−6^ ml cells^-1^ h^−1^ for *V* = 0.01 ml), although this argument has limits since plasmids cannot conjugate arbitrarily fast (indeed, Andrup and Andersen [1999] find that the number of transconjugants per donor saturates as the density of recipient cells increases). In our simulations, we consider population sizes of up to 10^6^ cells. Densities of 10^8^ CFU mL^−1^ are reached in some infections [Bingen et al., 1990, König et al., 1998] and a volume of *V* = 0.01 mL would thus not imply unrealistic densities even outside the lab. Finally, with a slower time scale, e.g. an intrinsic death rate of 1*/*10 h^−1^, our considered range of transfer coefficients would be between 10^−12^ and 10^−7^ ml cells^-1^ h^−1^. Overall, for a realistic model parameterization, there is a need for a better quantification of transfer rates along with other parameters such as population densities and birth and death rates in nature.

Throughout the paper, we made the assumption that the transfer coefficients of the resident and the rescue plasmid are identical within the focal population *β*_*r*_ = *β*_*R*_ = *β*. This is likely the case if the rescue plasmid is a mutated version of the resident plasmid that has been generated either in the focal population (Scenario A) or in an external population (Scenario B). However, when the resident and the rescue plasmid are compatible (Scenario C), this assumption is less realistic, because it implies either that there is a rather specific evolutionary relationship between the plasmids or that the transfer coefficient is fully determined by properties of the bacterial host and ecological factors rather than properties of the plasmid. Since in real life, the rate of conjugation is determined by both plasmid-related and plasmid-unrelated properties [Johnsen and Kroer, 2007], we analyze in the Supplementary Information a simple scenario in which the resident and rescue plasmid transfer rates are linearly correlated: *β*_*R*_ = *γβ*_*r*_ (section S3). We show that when the proportionality constant *γ* is large, negative effects of higher transfer rates can become dominant even in Scenario A (Figure S4). We furthermore also considered rescue if the transfer coefficients of rescue and resident plasmids are independent from each other. In that case, as expected, higher transfer rates of the rescue plasmid are always positive. Higher transfer coefficients of the resident plasmid, in contrast, can increase or decrease the rescue probability. Most interestingly, in Scenario A, the effect of an increase in the resident transfer rate depends on that of the rescue plasmid – if *β*_*R*_ is low, the effect is positive, if *β*_*R*_ is high, it is negative.

We assumed throughout the manuscript that the immigration rate *m* is constant, but ecological interactions between bacterial populations are common. In reality, the decline or rescue of the focal population could affect the external population size via competition, predation, or facilitation and thus the rate *m* at which inter-specific transfers occur [Bottery, 2022]. We further assumed that the rescue plasmid can appear in the focal population only after the environmental change. However rescue can also occur from alleles/genes that were already present in the population, i.e. from the standing genetic variation [Orr and Betancourt, 2001, Hermisson and Pennings, 2005]. It would be interesting to see how horizontal gene transfer affects rescue under these circumstances. More generally, a systematic study of the connection between horizontal transfer and standing genetic variation is to our knowledge missing in the literature.

Another potential extension of our model would be to take into account that plasmids are often present in multiple copies in the same host cell [Friehs, 2004]. Even though typically large plasmids with a single or a few copies can conjugate and small plasmids with many copies cannot conjugate, the latter can often still be mobilized during conjugation of the former [Smillie et al., 2010, Dionisio et al., 2019]. If the rescue plasmid itself is not conjugative but mobilizable by another plasmid, one would need to model the joint dynamics of both plasmids. Non-conjugative plasmids can moreover be horizontally transferred via transformation, transduction, or in membrane vesicles [Rodríguez-Rubio et al., 2020, Tran and Boedicker, 2019]. Santer and Uecker [2020] studied evolutionary rescue through mutations on non-transmissible multicopy plasmids and showed that the probability of rescue is influenced by the dominance relationship between rescue and resident plasmids. To our knowledge, Bichsel et al. [2010] is the only theoretical work on multicopy mobile DNA but the establishment probability of a new gene is determined in the absence of a resident gene. In other words, they do not consider competition at the plasmid-level nor dominance relationships. Such a theoretical study remains to be done. Transfer of a single mutant plasmid copy would be a jackpot event since the recipient cell is homozygous for the mutant plasmid. This could give the establishment probability of the mutation a boost, which could be particularly relevant for recessive mutations.

The conjugation rate is a fixed parameter in our models and cannot evolve. Moreover, selection for a lower or higher transfer rate would only be indirect through a higher chance of survival of the plasmid or bacteria (whoever is controlling the rate of transfer). It is nevertheless interesting to briefly reconsider our results in the context of the evolution of conjugation rates. The rate of transfer can be controlled by the plasmid or the bacteria (either the donor or the recipient cell), and interests of the two agents are not always aligned, as shown by Sheppard et al. [2021]. In our study, we focused on survival of the bacterial population. Survival of the bacteria does not always entail survival of the resident plasmid though: especially, if rescue occurs through an incompatible immigrant plasmid (Scenario B), the resident plasmid is doomed to extinction even if the bacterial population survives. In Scenario A, both are rescued together, and in Scenario C, the resident plasmid can hitchhike to rescue. For the resident plasmid, it is therefore almost always optimal to transfer as much as possible. For the bacteria, in contrast, in can be favorable to enforce a reduced transfer rate to maintain plasmid-free recipient cells. One could speculate that keeping the option to receive plasmids is overall more important for the bacterial population than the benefit gained through a higher chance of rescue by *de novo* mutations on plasmids. Our results show that even if transfer rates were selected to maximise survival of the bacteria or the plasmid, there would not be a transfer rate that is optimal under all circumstances and for both agents.

Which genes are typically carried on plasmids or on the chromosome – and why – is a focal question about the evolutionary biology of plasmids. For example, genes located on plasmids that are beneficial for the bacterial host often code for functions that are useful in some restricted temporal or spatial niches, such as heavy metal tolerance and antibiotic resistance [Rankin et al., 2011, Harrison and Brockhurst, 2012, Carroll and Wong, 2018]. The reason could be that these genes, if on plasmids, can be acquired though horizontal gene transfer when needed and lost again when only an unnecessary burden [Eberhard, 1990, Bergstrom et al., 2000]. Is rescue more likely if the relevant gene is on the plasmid or on the chromosome? Within our model, the comparison is most meaningful for rescue by *de novo* mutations. Tazzyman and Bonhoeffer [2014] found that, depending on the transfer coefficient and the fraction of resident plasmid cells, rescue can be more or less likely on a plasmid than on a chromosome. When the dynamics of resident plasmid cells are taken into account (Model II), we show in the Supplementary Information that rescue is actually never more likely to occur on a plasmid than on the chromosome (SI section S5, Figure S8). Yet, this might change if the plasmid copy number is greater than one as has been found for non-transmissible plasmids [Santer and Uecker, 2020].

Our model was designed to study adaptation on a conjugative plasmid. However, it could be extended to the other modes of horizontal transfer, i.e. transformation, transduction, and membrane vesicles [Chen and Dubnau, 2004, Brüssow et al., 2004, Domingues and Nielsen, 2017], as long as plasmid transfer follows a mass-action model with donor and recipient cells (but note that for transformation, mass action kinetics might not capture transfer well since the rate of take-up depends on the amount of DNA in the environment as pointed out by Leclerc et al. [2019]). The main difference would lie in the mechanism of plasmid incompatibility. Indeed, we have assumed that a conjugative plasmid cannot enter a cell that already bears an incompatible plasmid due to surface exclusion, but this mechanism is restricted to conjugation [Garcillán-Barcia and de la Cruz, 2008]. In the other modes of horizontal transfer, incompatibility is the result of unstable coexistence of both plasmid types in the same host cell [Kittell and Helinski, 1993]. Either the new or the already-present plasmid will be lost in subsequent replication. The rescue dynamics would be different then, since resident plasmids now impair the growth of rescue plasmid cells by potentially converting them into resident plasmid cells.

In conclusion, we provided a detailed study of evolutionary rescue on conjugative plasmids based on Tazzyman and Bonhoeffer [2014] with a focus on the interplay and feedbacks between horizontal gene transfer and the dynamics of competition at the plasmid level. Although our model includes only the most fundamental processes, the results reveal that the effects of plasmid transfer on adaptation are surprisingly complex. There is no conjugation rate that is universally optimal. In several different scenarios, higher levels of horizontal transfer do not necessarily promote adaptation on conjugative plasmids due to competition between resident and rescue plasmids for plasmid-free cells. Higher rates of horizontal transfer can be deleterious even when the plasmids are compatible, thus in the absence of plasmid competition, because the presence of costly, highly infectious resident plasmids is a burden that can reduce the total resident population size. We hope that our systematic approach helps to understand the various effects of plasmid transfer on rescue and that our study will form the starting point for future models that take more ecological and genetic complexities into account.

## Data accessibility

The source code for numerical and stochastic simulation results is available on the first author’s GitHub repository: https://github.com/fgeoffroy/rescue_plasmid.

## Acknowledgments

We thank the members of The Rescue Team at the Max Planck Institute for Evolutionary Biology and Sally Otto for helpful discussions. We are grateful to several anonymous reviewers for valuable comments on the manuscript. This work was funded by the Deutsche Forschungsgemeinschaft (DFG, German Research Foundation) – project numbers 418432175 (https://gepris.dfg.de/gepris/projekt/418432175); 400993799 (project 3 within the Research Training Group 2501 “Translational Evolutionary Research”, https://gepris.dfg.de/gepris/projekt/400993799). Figure 1 was created with BioRender.com.

## Authorship contributions

F.G.: conceptualization, methodology, formal analysis, software, visualization, writing—original draft H.U.: conceptualization, methodology, visualization, supervision, writing—review and editing

## A Fraction of resident plasmid cells

In this section, we provide the mathematical analysis for the dynamics of the fraction of resident plasmid cells *φ*(*t*) in Models II and III. The dynamics of plasmid-free cells *N*_*F*_ (*t*) and resident plasmid cells *N*_*r*_(*t*) after the environmental change is given by (see system (3) in the main text):

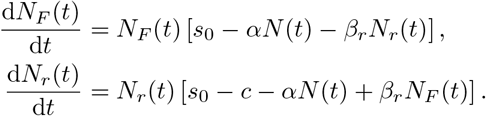

Given *N* (*t*) = *N*_*F*_ (*t*) + *N*_*r*_(*t*) and *φ*(*t*) = *N*_*r*_(*t*)*/N* (*t*), applying the quotient rule gives the equivalent system of ordinary differential equations:

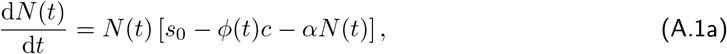

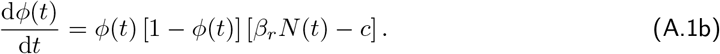

If *c* = 0, we have 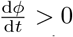 except for the trivial situations *φ*(*t*) = 0, *φ*(*t*) = 1, or *β*_*r*_ = 0. Hence, the fraction of resident plasmid cells *φ*(*t*) is always increasing. Furthermore, since *s*_0_ *<* 0, the total resident population is going to extinction, lim_*t→∞*_ *N* (*t*) = 0, and thus 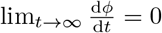. Namely the fraction of resident plasmid cells *φ*(*t*) converges as the resident population is going to extinction (see Figure 4a in the main text).

If *c >* 0, we have 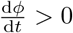 if 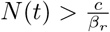 and 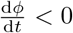 if 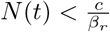. The fraction of resident plasmid cells might thus increase at first when enough plasmid-free cells are present so that the positive effect of horizontal transfer is larger than the negative effect of plasmid cost. However, since lim_*t→∞*_ *N* (*t*) = 0, the effect of plasmid costs will eventually always become dominant such that lim_*t→∞*_ *φ*(*t*) = 0, i.e. the fraction of resident plasmid cells is going to zero as the resident population is going to extinction (see Figure 4a in the main text). In the next section, we show that, if we determine the initial numbers of plasmid-free and resident plasmid-bearing cells from the pre-rescue dynamics, the fraction of resident plasmid cells always decreases.

### B Pre-rescue dynamics

In this section, we provide the mathematical analysis for the equilibrium of the pre-rescue dynamics in Model III. The dynamics of plasmid-free cells *N*_*F*_ and resident plasmid cells *N*_*r*_ before the environment changes is given by (see system (4) in the main text):

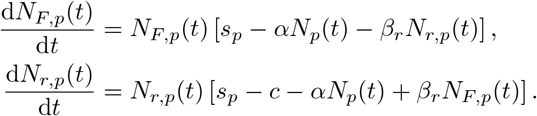

We assume that both types can grow in the absence of the other type, i.e. *s*_*p*_ *>* 0 and *s*_*p*_ − *c >* 0. There are four possible equilibria 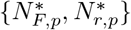:

- {0, 0} the ‘null equilibrium’
- 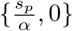 the ‘plasmid-free equilibrium’
- 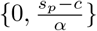 the ‘resident plasmid equilibrium’
- 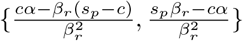 the ‘mixed equilibrium’

We discard the ‘null equilibrium’ since it excludes the possibility of a rescue scenario. An equilibrium is stable if the real parts of both eigenvalues of the following Jacobian matrix evaluated at this equilibrium are negative:

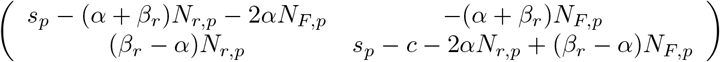

Three cases depending on the value of *β*_*r*_ can be distinguished.

- If 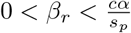, only the ‘plasmid-free equilibrium’ has positive population size values and is stable (negative Jacobian eigenvalues).
- If 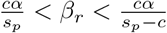, only the ‘mixed equilibrium’ has positive population size values and is stable.
- If 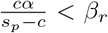, only the ‘resident plasmid equilibrium’ has positive population size values and is stable.

*Effect of β*_*r*_ *on the ‘mixed equilibrium’*

When both cell types coexist at the equilibrium, it holds that 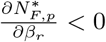, i.e. more frequent horizontal transfer decreases the number of plasmid-free cells at the equilibrium. However, the effect of *β*_*r*_ on the the number of resident plasmid cells at the equilibrium can be non-monotonous:

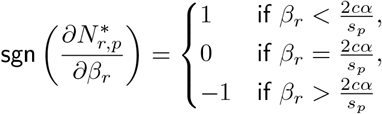

with the sign function 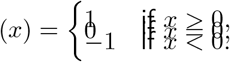,

Note that, since the value of *β*_*r*_ is bounded for the mixed equilibrium, the third condition is only possible if 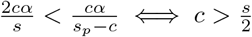. Thus, if both the transfer coefficient *β*_*r*_ and the plasmid cost *c* are large enough, more frequent horizontal transfers can actually decrease the number of resident plasmid cells at the equilibrium.

For the total number of cells, we have:

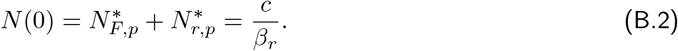

The fraction of resident plasmid cells increases with *β*_*r*_:

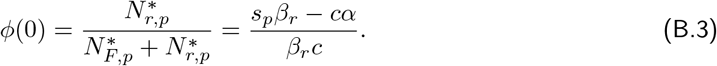

Figure B.1a shows the fraction of resident plasmid cells at equilibrium.

Lastly, we can show that, if the pre-rescue dynamics leads to a mixed population at the equilibrium, then the fraction of resident plasmid cells *φ*(*t*) will always decrease after the environment has changed (when we allow *φ* to vary). At time *t* = 0, using 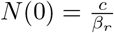 in Eq. (A.1b), we have 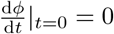. Since the population size *N* (*t*) is decreasing, we have 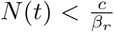 for *t >* 0 and thus 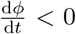 for all *t >* 0.

**Figure B.1:**
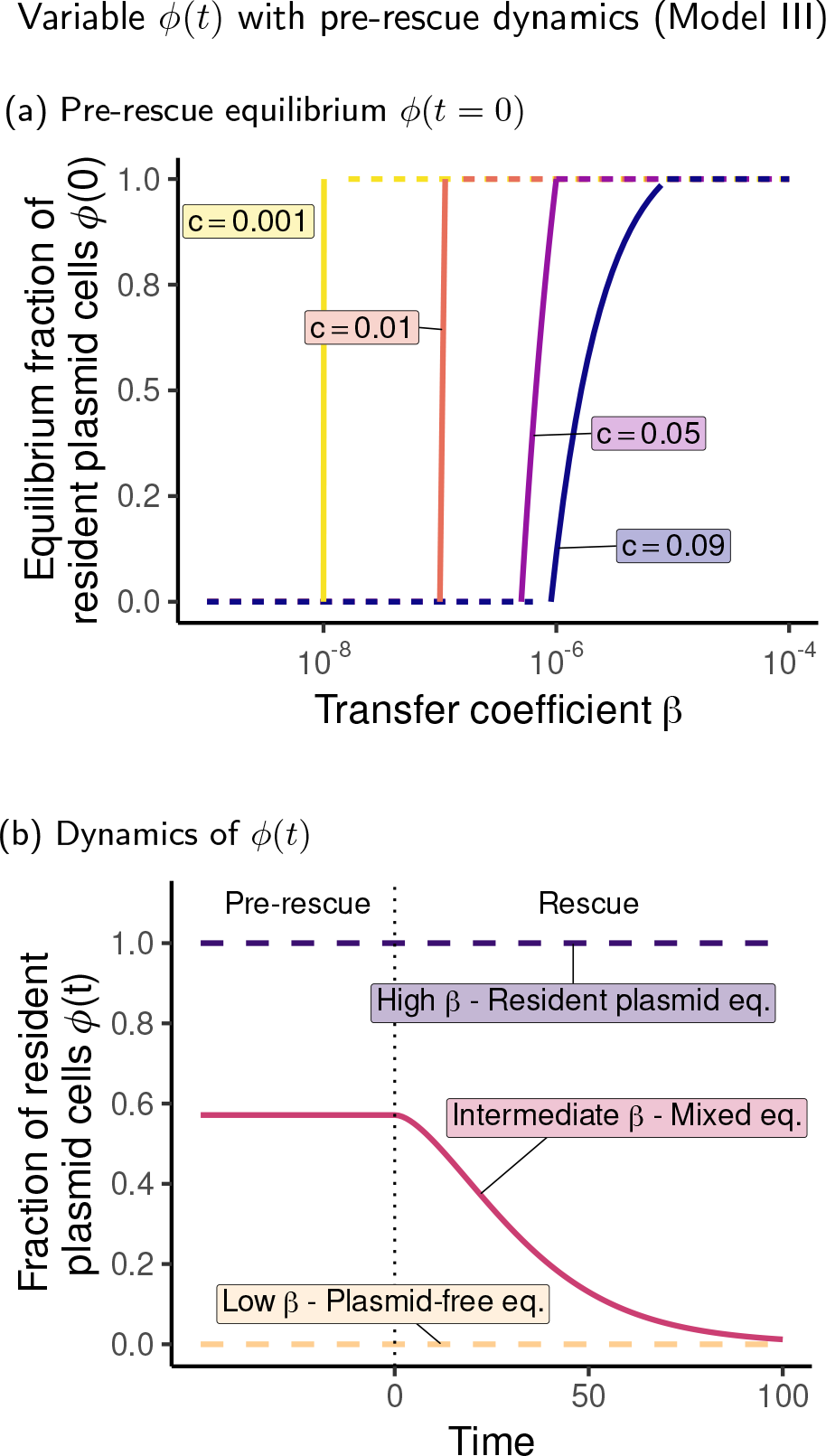
(a) The fraction of resident plasmid cells at equilibrium before the environmental change 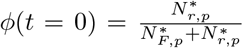. (b) The fraction of resident plasmid cells before the environmental change (‘Pre-rescue’) and after the environmental change (‘Rescue’) for three transfer coefficient values: *β* ∈ {5 *·* 10^−7^; 7 *·* 10^−7^; 10^−6^}. The same parameters as in Figures 3, 5 and 6 in the main text were used.

### C Dependence of the immigration rate *m* on the transfer coefficient *β*

In the main text, we made the assumption that the rate *m* at which rescue plasmids are being transferred from an external population (Scenarios B and C) is independent of the rate *β*_*R*_ at which they are then being transferred within the focal population. In this section, we assume that the immigration rate is proportional to the transfer coefficient, i.e. *m* = *κβ*_*R*_. For the sake of simplicity, we assume that the horizontal transfer coefficients of the resident and the rescue plasmid are identical (*β*_*r*_ = *β*_*R*_ = *β*).

We are interested in examining how this new assumption changes the effect of the transfer coefficient *β* on the rescue probability. Indeed, now, higher values of the transfer coefficient *β* imply an increase of the immigration rate *m*, but not necessarily an increase of the net rate of appearance *λ*(*t*) which equals *mN*_*F*_ (*t*) in Scenario B and *m*(*N*_*F*_ (*t*) + *N*_*r*_(*t*)) in Scenario C.

In Model I, the transfer coefficient *β* affects neither *N*_*F*_ (*t*) nor *N*_*r*_(*t*) since transfers of resident plasmids are not explicitly modelled. Accordingly, higher values of *β* always result in higher chances of evolutionary rescue (results not shown). The only difference with the results in the main text is that the effect of *β* on rescue is now also mediated by its positive effect on the immigration rate.

Figure C.1 shows the rescue probability for Models II and III in Scenario B and C. When compared to the results presented in the main text (Figures 5b,c and Figures 6b,c, where the immigration rate *m* is an exogeneous constant), we observe that the transfer coefficient *β* has overall a larger positive effect on rescue, but an increase in *β* can still decrease the probability of rescue. This is especially true when the rescue plasmid is incompatible (Scenario B) and pre-rescue dynamics are taken into account (Model III); see Figure C.1c. In this case, large enough transfer coefficients *β* result in a resident population without plasmid-free cells, leading to a null net rate of appearance *λ*(*t*), irrespective of *m*. The transfer coefficient *β* can also still decrease the probability of rescue in Model II, but only for large *β* (see Figure C.1a). This indicates that the effect of the transfer coefficient *β* becomes more similar between Models I and II when we make the more realistic assumption that the immigration rate *m* depends on the transfer coefficient *β*.

We directly compare Models I and II in Figure C.2 for *m* = const. and for *m* = *κβ*. While under the more realistic assumption on *m*, Models I and II become more similar regarding the sign of the effect of *β* (whether is has a positive or a negative effect on the probability of rescue) in Scenario B, the model predictions differ more in absolute values for intermediate and high *β* values than for *m* = const. For Scenario C, the positive effect of *β* on the rescue probability in Models I and II is amplified for *m* = *κβ* compared to *m* = const. (Figure C.2b).

**Figure C.1:**
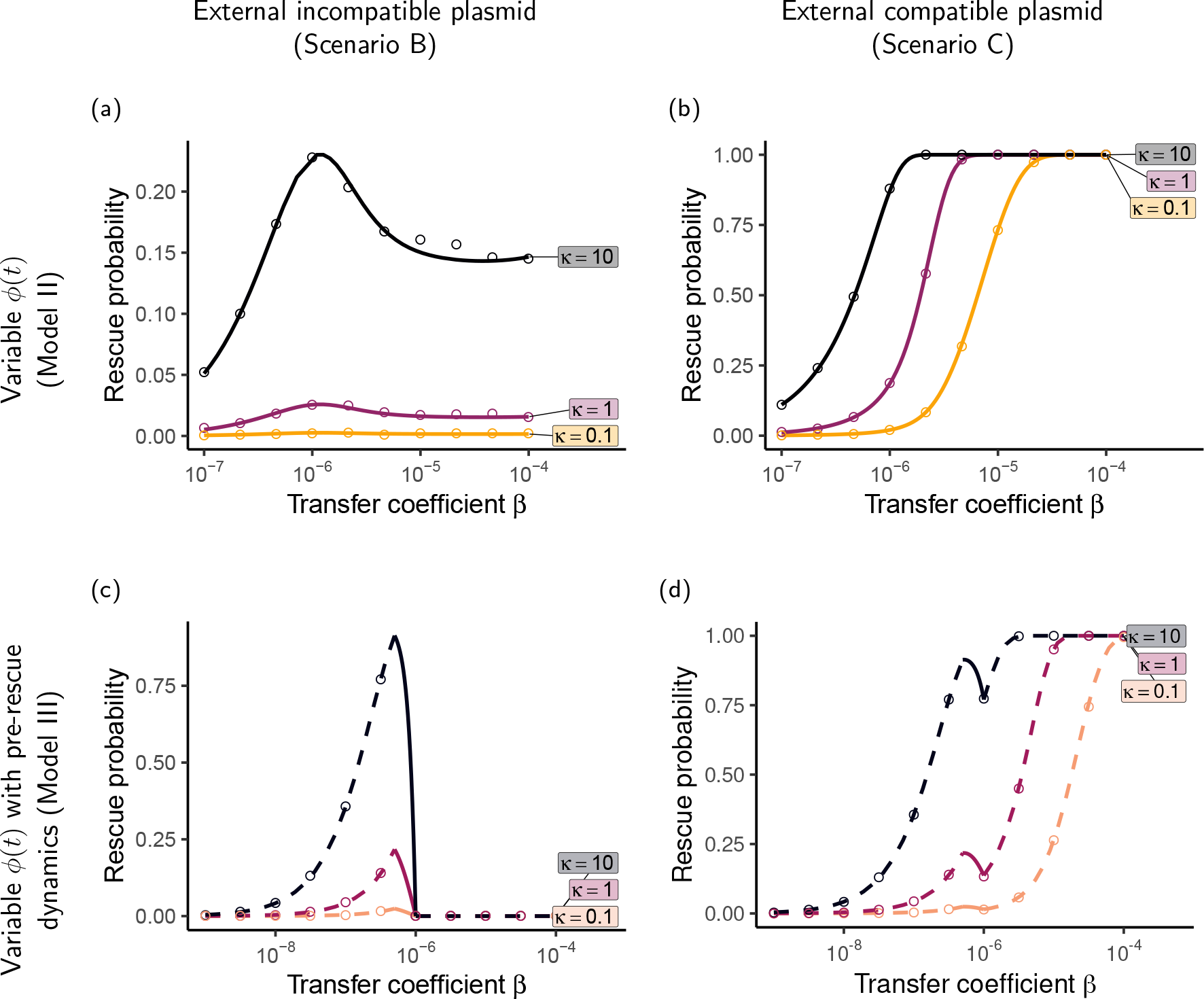
The probability of evolutionary rescue when the immigration rate of rescue plasmids depends linearly on the plasmid transfer rate (*m* = *κβ*). Parameters: (a)-(b) *φ*_0_ = 1*/*2 ; *c* = 0 ; (c)-(d) *s*_*p*_ = 1 ; *s*_*R*_ = 1 ; *c* = 0.5. All other parameters are the same as in Figures 3, 5, and 6 in the main text. Note that the range of the *x*-axis differs between Panels (a)-(b) and (c)-(d) and that the range of the *y*-axis varies across panels.

**Figure C.2:**
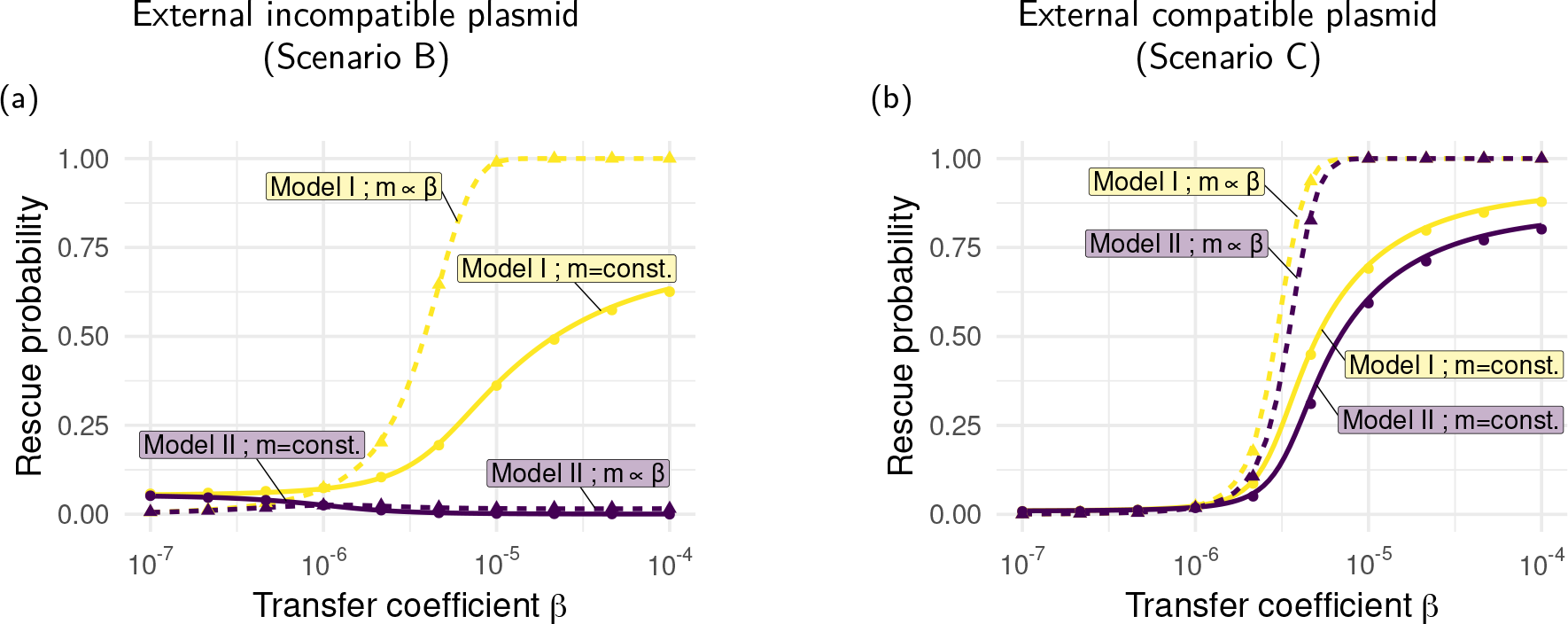
Comparison of Models I, and II with the additional assumption that the plasmid immigration rate *m* linearly depends on the transfer coefficient *β*, i.e. *m* = *κβ*. Parameters: *κ* = 1; *φ* = 0.5; (a) *c* = 0 ; (b) *c* = 0.09.

