## Supplementary Information for "Limits to evolutionary rescue by conjugative plasmids"

#### S1 Alternative model assumptions concerning $s$, $c$, and $\alpha$

In the main text, we assumed that the plasmid type (resident or rescue) affects the cell division rate but not the death rate and that plasmid costs and competition increase the death rate, while leaving the cell division rate unchanged. In this section, we show for Scenario A (rescue by *de novo* mutations) that alternative assumptions do not qualitatively change our main results.

In a first model variant, we make the assumption that the parameters $s_0$ and $s_R$ enter via the death rate rather than the replication rate, and set the replication rate (rather than the death rate) to $1$. Note that this modification also affects the rate $\lambda(t)$ at which rescue plasmids appear. We thus have:

$$
\begin{aligned}
b(t) &= 1 + \beta_R N_F(t),\\
d(t) &= 1 - s_R + c + \alpha N(t),\\
\lambda(t) &= u N_r(t).
\end{aligned}
$$

In a second model variant, we make the assumption that the plasmid cost $c$ affects the replication rate rather than the death rate, thus:

$$
\begin{aligned}
b(t) &= 1 + s_R - c + \beta_R N_F(t),\\
d(t) &= 1 + \alpha N(t),\\
\lambda(t) &= u(1 + s_0 - c) N_r(t).
\end{aligned}
$$

In a third model variant, we make the assumption that competition, modeled with the parameter $\alpha$, affects the replication rate rather than the death rate, thus:

$$
\begin{aligned}
b(t) &= 1 + s_R - \alpha N(t) + \beta_R N_F(t),\\
d(t) &= 1 + c,\\
\lambda(t) &= u(1 + s_0 - \alpha N(t)) N_r(t).
\end{aligned}
$$

Note that there are additional constraints on the parameters to prevent the replication and death rates from becoming negative. Results for the probability of rescue under the three model variants are shown in Figure S1. Overall, we see that the above changes in the model assumptions do not change the qualitative trends that we observe in the main text in Figures 3, 5 and 6. In model variants with lower replication rates, the overall rescue probability is reduced. For constant $b$ and $d$ and $b - d$, the establishment probability is given by $p_{\text{est}} = \frac{s}{b}$ and thus increases with decreasing $b$. However the rate $\lambda(t)$ at which rescue plasmids appear decreases with $b$, which seems to be the stronger effect.

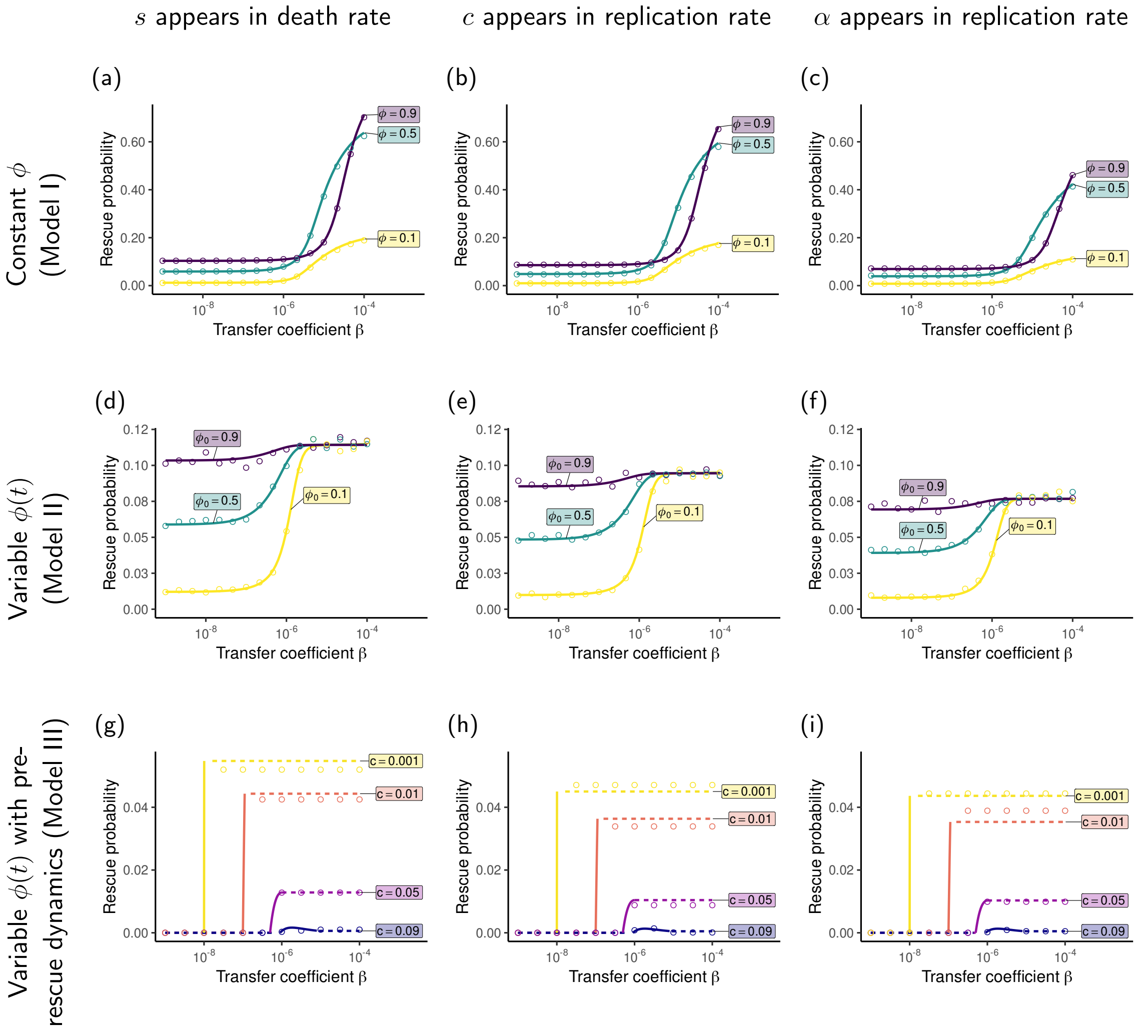


Figure S1: The probability of evolutionary rescue by *de novo* mutations (Model A) under alternative modeling assumptions regarding the parameters $s$, $c$, and $\alpha$. Alternative implementations of the effects of $s$, $c$ and $\alpha$ do not qualitatively change the results. The same parameters as in Figures 3, 5, and 6 in the main text were used.

### S2 Further comparison between Models I and II

In the main text, we put results from Models I and II next to each other for which the constant $\Phi$ in Model I was the initial value $\Phi_0$ in Model II. I.e., we asked about the probability of evolutionary rescue of a population when we know the fraction of resident plasmid cells at time $0$, assuming either that the fraction remains constant or changes over time. In this section, we make a different comparison between Models I and II, setting $\Phi$ in Model I to the average (rather than the initial) fraction in Model II. More precisely, we first compute the average fraction of resident plasmid cells in Model II between $t = 0$ and $T_{\text{res}}$, the time at which the resident population size is reduced to a single cell: $E(\phi(t)) = \int_0^{T_{\text{res}}} \phi(t)\,\mathrm{d}t$. Averaged values of $\phi$ are shown in Figure S2 for multiple values of the initial fraction $\phi_0 = \phi(t = 0)$ and of the transfer coefficient $\beta$. Then, we use the previously computed values of $E(\phi(t))$ from Model II as constant $\phi$ in Model I.

Figure S3 displays the corresponding rescue probabilities only for Scenarios A and B, since $\phi$ does not have any influence on rescue in Scenario C where it is constant (see Figure 6a in the main text). These results are qualitatively and quantitatively similar to those obtained in Model II (see Figures 3b and 5b in the main text); the rescue probability can saturate or even decrease with higher transfer coefficient values. This indicates that results of Model II are mainly driven by a change in the fraction of resident plasmids to a higher average value, rather than by the specific dynamics.

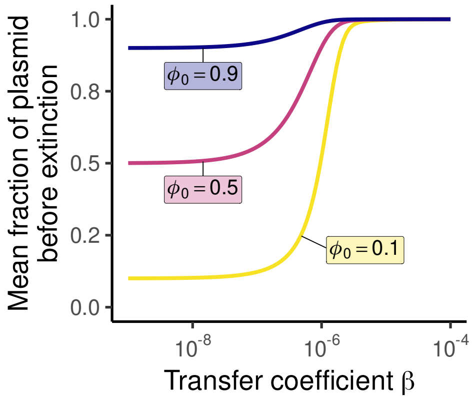


Figure S2: The average fraction of resident plasmid cells $E(\phi(t))$ for Model II between $t = 0$ and $T_{\text{res}}$, the time at which the resident population size is reduced to a single cell. The same parameters as in Figures 3 and 5 in the main text were used.

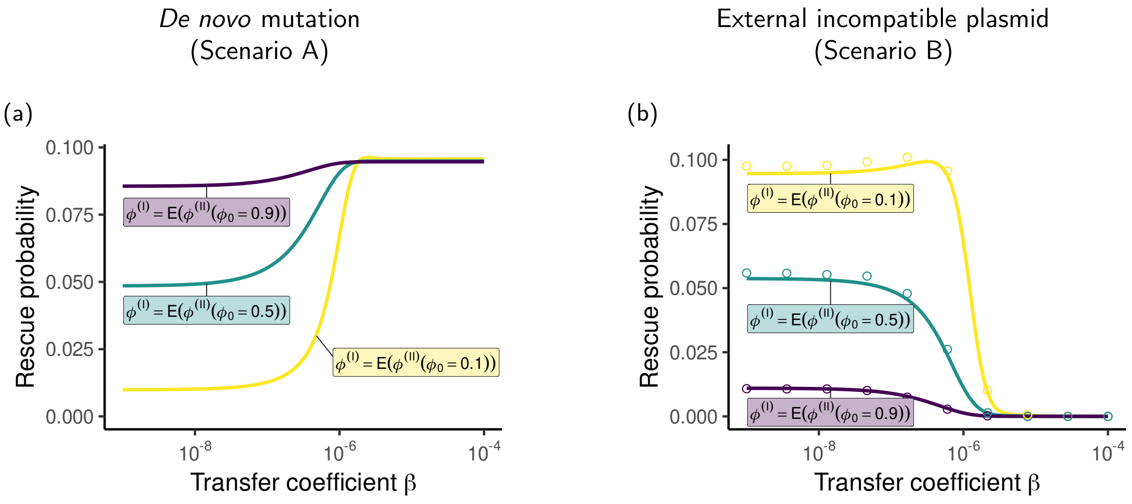


Figure S3: The probability of evolutionary rescue in Model I when the constant fraction $\phi$ of resident plasmid cells is chosen as the average value of the variable $\phi(t)$ in Model II for three initial values $\phi(t=0) \in \{0.1, 0.5, 0.9\}$. The average $E(\phi(t))$ is computed between $t=0$ and $T_{\text{res}}$, the time at which the resident population size is reduced to a single cell (see Figure S2). The same parameters as in Figures 3 and 5 in the main text were used.

#### S3 Plasmid-type specific transfer coefficients

In the main text, we assumed that the horizontal transfer coefficients of the resident and the rescue plasmid are identical, i.e. $\beta_r = \beta_R = \beta$. In this section, we instead assume (i) that the transfer coefficients are proportional to each other, $\beta_R = \gamma\beta_r$, or (ii) that the transfer coefficients are completely uncorrelated. The idea behind assumption (i) is that host control of conjugation rates could affect both plasmids.

Figure S4 shows the rescue probability for our different scenarios (excluding Model I, for which transfers of resident plasmids are not explicitly modelled) when the transfer coefficients are proportional to each other. By comparing the curves $\gamma = 1$ (which correspond to the results in the main text, Figures 3b-c, 5b-c, and 6b-c) to the others, one can see that many of our results are qualitatively robust to this change of assumption. However, for Models A and B, and for large values of the proportionality constant ($\gamma = 10$, Figures S4a, S4b and S4d), the rescue probability becomes non-monotonic in the transfer coefficient $\beta_r$. This can be intuitively understood by considering the two-fold effect of increasing $\beta_r$ on the rescue probability in different regimes:

- For low values of $\beta_r$, the competition between the two plasmid types is low because many plasmid-free cells are present. A small increase of $\beta_r$ would lead to a much larger increase of $\beta_R$ because the proportionality constant is high. As a consequence, the benefit of increasing the effective fitness of rescue plasmid cells would be much larger than the cost of plasmid competition Figures S4a and S4d) as well as the cost of reducing the rate of appearance of an incompatible rescue plasmid from an external population (Figure S4b).
- For high values of $\beta_r$, the effect of plasmid competition is so strong that increasing the transfer coefficient of the rescue plasmid is pointless because there are almost no plasmid-free cells that they can 'infect'. In that case, the negative effects of increasing $\beta_r$ are larger than the benefits and thus, the rescue probability decreases.

Figure S5 shows the probability of evolutionary rescue in the same scenarios, assuming that the transfer coefficients $\beta_r$ and $\beta_R$ are completely decorrelated. In that case, it is easier to decompose the effect of each of them. High transfer coefficients of the rescue plasmid $\beta_R$ always increase the probability of evolutionary rescue. High transfer coefficients of the resident plasmid $\beta_r$ mostly reduce the chances of rescue, which is due to plasmid competition but also to a negative effect on the initial population size if pre-rescue dynamics are taken into account (note that costs are high in Figure S5d-f). Only if rescue relies on a *de novo* mutation on the resident plasmid (Scenario A, Figures S5a and S5d) such that resident plasmid cells are required for the appearance of rescue plasmids, can it sometimes be beneficial for the population to have a resident plasmid that can transfer quickly. Note however, that this is only the case if (i) the transfer coefficient of rescue plasmids $\beta_R$ is low enough such that horizontal transfer does not contribute much to its effective fitness (Figure S5a), or (ii) when pre-rescue dynamics are taken into account, in which case the resident plasmid is absent if its transfer coefficient is too low (Figure S5d). Note that even if the two plasmids are compatible, a high transfer coefficient of the resident plasmid can impede rescue through its negative effect on the total population size (Figure S5f).

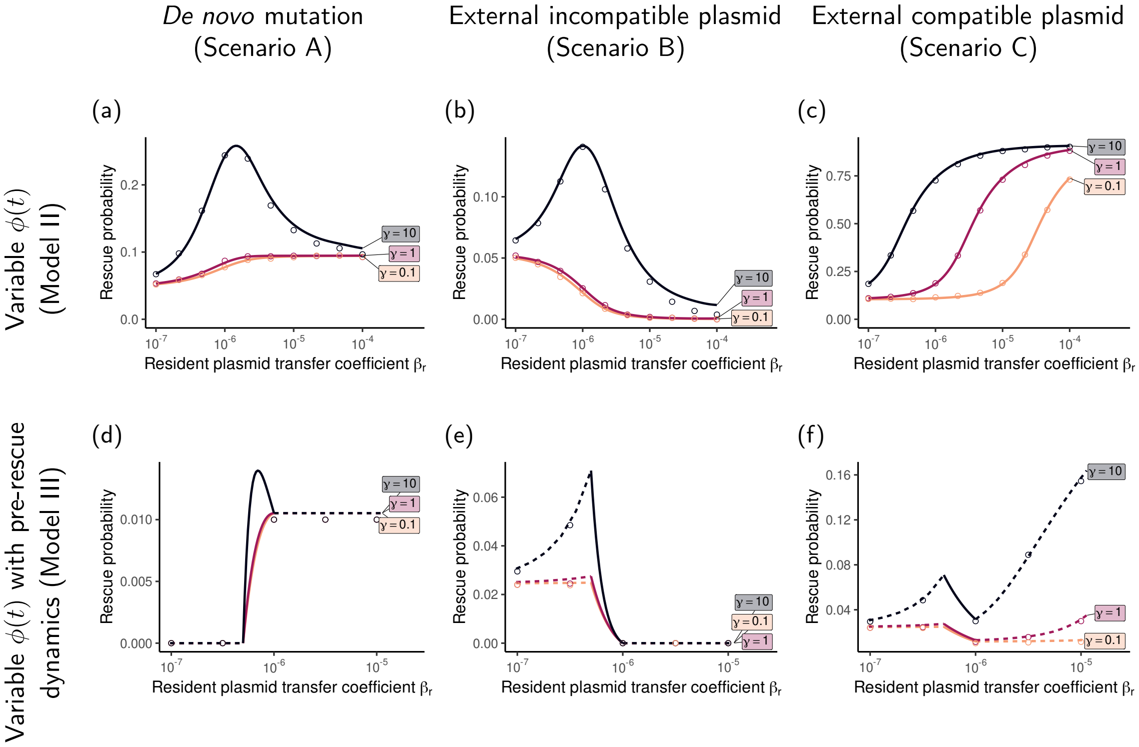


Figure S4: The probability of evolutionary rescue when the transfer coefficients of the resident plasmid ($\beta_r$) and the rescue plasmid ($\beta_R$) are proportional to each other ($\beta_R = \gamma \times \beta_r$). In Panels (a)-(c), the initial fraction of resident plasmid cells is $\phi_0 = 1/2$ and plasmids impose no cost on their hosts ($c = 0$). In Panels (d)-(f), the plasmid cost is $c = 0.5$. All other parameters are the same as in Figures 3, 5, and 6 in the main text. Note that the range of the $x$-axis differs between Panels (a)-(c) and (d)-(f) and that the range of the $y$-axis varies across panels.

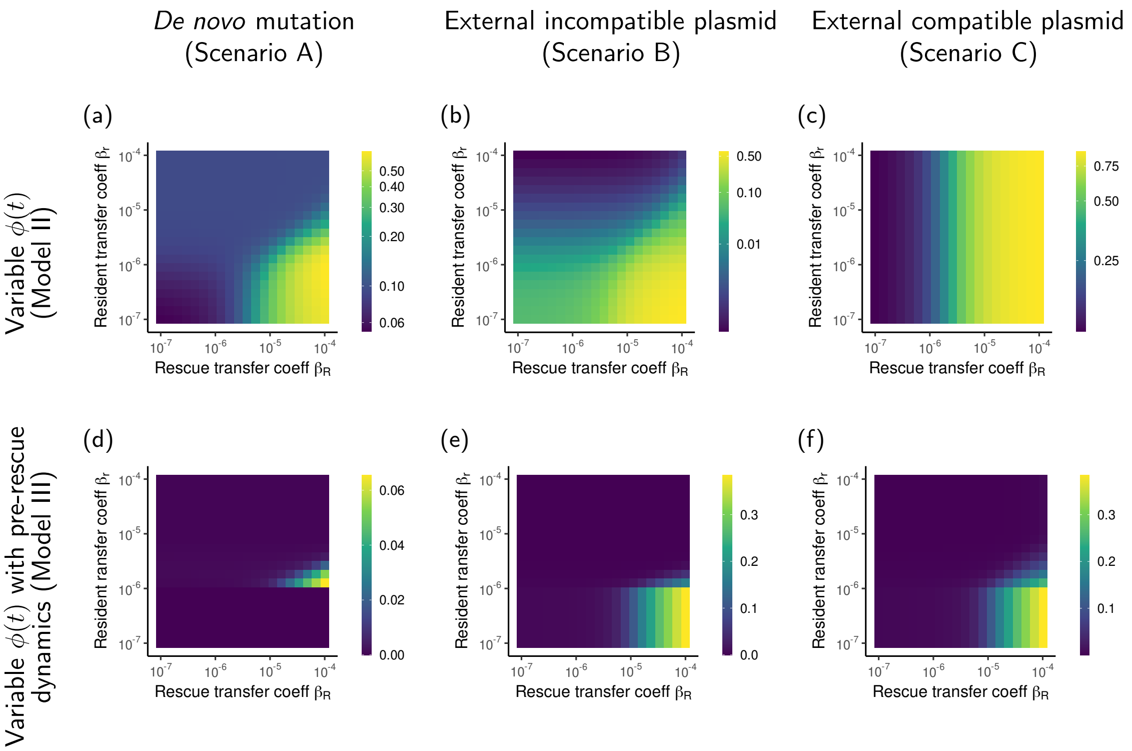


Figure S5: The probability of evolutionary rescue when the transfer coefficients of the resident plasmid ($\beta_r$) and the rescue plasmid ($\beta_R$) are independent from each other. In Panels (a)-(c), the initial fraction of resident plasmid cells is $\phi_0 = 1/2$ and plasmids impose no cost on their hosts ($c = 0$). In Panels (d)-(f), the plasmid cost is $c = 0.09$. All other parameters are the same as in Figures 3, 5, and 6 in the main text. Note that the color scales are different between plots. In particular, a logarithmic scale is used in Panels (a)-(c) and a linear scale in Panels (d)-(f).

### S4 The effect of segregational loss $\sigma$ on the probability of evolutionary rescue

In this section, we assume that with probability $\sigma$, one of the daughter cell does not receive a copy of the plasmid at replication of a plasmid-bearing cell ('segregational loss'). For simplicity, we do not explicitly consider cells with two copies of a plasmid, but we rather assume that they behave like cells with a single copy.

The replication rate of a rescue plasmid cell becomes:

$$\begin{aligned} \text{'birth rate' (incompatible rescue plasmid):} \quad b^{(i)}(t) &= (1+s_R)(1-\sigma) + \beta_R N_F(t), \\ \text{'birth rate' (compatible rescue plasmid):} \quad b^{(c)}(t) &= (1+s_R)(1-\sigma) + \beta_R (N_F(t) + N_r(t)). \end{aligned}$$

##### Model I - Constant $\phi$

When the fraction of resident plasmid cells $\phi$ is held constant, segregational loss of resident plasmids is not explicitly considered, but rather the value of $\phi$ is considered to be the result of horizontal transfers, plasmid cost and segregation Tazzyman and Bonhoeffer [2014]. Hence, the introduction of segregational loss simply reduces the establishment probability compared to the results shown in the main text. As a consequence, the probability of evolutionary rescue is also reduced (results not shown).

##### Model II - Variable $\phi(t)$

When we relax the assumption of a constant fraction of resident plasmid cells $\phi$, the dynamics of the resident population becomes:

$$\frac{\mathrm{d}N_F}{\mathrm{d}t}(t) = N_F(t)\left[(1+s_0) - 1 - \alpha N(t) - \beta_r N_r(t)\right] + N_r(t)(1+s_0)\sigma, \tag{S1a}$$

$$\frac{\mathrm{d}N_r}{\mathrm{d}t}(t) = N_r(t)\left[(1+s_0)(1-\sigma) - (1+c) - \alpha N(t) + \beta_r N_F(t)\right]. \tag{S1b}$$

With $N(t) = N_F(t) + N_r(t)$ and $\phi(t) = N_r(t)/N(t)$, the system is equivalent to

$$\frac{\mathrm{d}N}{\mathrm{d}t}(t) = N(t)\left[s_0 - \phi(t)c - \alpha N(t)\right], \tag{S2a}$$

$$\frac{\mathrm{d}\phi}{\mathrm{d}t}(t) = \phi(t)\left[(1-\phi(t))\left(\beta_r N(t) - c\right) - \sigma(1+s_0)\right]. \tag{S2b}$$

We can use the same reasoning as in Appendix A in the main document: the total resident population is going to extinction eventually $\lim_{t\to\infty} N(t) = 0$. Therefore, $\frac{\mathrm{d}\phi}{\mathrm{d}t}(t)$ is negative once the population size has been sufficiently reduced, even in the absence of plasmid costs ($c = 0$).

Figures S6a, S6b and S6c show that plasmid segregation does not qualitatively change the main trends in the effect of the transfer coefficient $\beta$, namely that higher values of $\beta$ increase the rescue probability for Scenarios A and C but not for Scenario B (see Figures 3b, 5b, and 6b in the main text). Interestingly, even though plasmid segregation overall decreases the probability of rescue, it can have a positive effect when the rescue plasmid comes from an external population and is incompatible with the resident plasmid (Figure S6b). In that case, plasmid segregation generates plasmid-free cells which can later receive the rescue plasmid from the external population, leading to rescue. However, for larger probabilities of segregational loss, the negative effects on the effective fitness of rescue plasmid cells becomes dominant and the effect is reversed (Figure S7a).

##### Model III - Variable $\phi(t)$ with pre-rescue dynamics

With plasmid segregation, the dynamics of plasmid-free cells $N_F$ and resident plasmid cells $N_r$ before the environment changes are given by:

$$\frac{\mathrm{d}N_{F,p}}{\mathrm{d}t}(t) = N_{F,p}(t)\left[(1+s_p) - 1 - \alpha N_p(t) - \beta_r N_{r,p}(t)\right] + N_{r,p}(t)(1+s_0)\sigma, \tag{S3a}$$

$$\frac{\mathrm{d}N_{r,p}}{\mathrm{d}t}(t) = N_{r,p}(t)\left[(1+s_p)(1-\sigma) - (1+c) - \alpha N_p(t) + \beta_r N_{F,p}(t)\right]. \tag{S3b}$$

Contrary to the case without plasmid segregation (see Appendix B in the main document), there cannot be an equilibrium with only resident plasmid cells because plasmid-free cells are constantly generated by segregation (assuming $\sigma > 0$). Thus, disregarding the trivial 'null equilibrium', there are only two possible equilibria:

- $\{\frac{s_p}{\alpha}, 0\}$ the 'plasmid-free equilibrium'
- $\{N^*_{F,p}, N^*_{r,p}\}$ the 'mixed equilibrium' with

$$N^*_{F,p} = \frac{c(\alpha+\beta_r) + \sqrt{4\alpha\beta_r c\sigma(s_p+1) + (\beta_r s_p - c(\alpha+\beta_r))^2} + 2\beta_r\sigma(s_p+1) + \beta_r(-s_p)}{2\beta_r^2},$$

$$N^*_{r,p} = \frac{-c\left(\alpha^2+\beta_r^2\right) - \alpha\sqrt{4\alpha\beta_r c\sigma(s_p+1) + (\beta_r s_p - c(\alpha+\beta_r))^2} + \beta_r\sqrt{4\alpha\beta_r c\sigma(s_p+1) + (\beta_r s_p - c(\alpha+\beta_r))^2} - 2\alpha\beta_r\sigma(s_p+1) + \beta_r s_p(\alpha+\beta_r)}{2\alpha\beta_r^2}.$$

Regarding the stability of those equilibria, two cases depending on the value of $\beta_r$ can be distinguished.

- If $0 < \beta_r < \alpha\frac{c+\sigma(1+s_p)}{s_p}$, only the 'plasmid-free equilibrium' has positive population size values and is stable.
- If $\alpha\frac{c+\sigma(1+s_p)}{s_p} < \beta_r$, only the 'mixed equilibrium' has positive population size values and is stable.

Again, Figures S6d, S6e and S6f show that plasmid segregation does not qualitatively change the main in the effect of $\beta$ (see Figures 3, 5 and 6 in the main text). As for Model II, plasmid segregation has a positive effect on evolutionary rescue if rescue relies on the immigration of an external incompatible rescue plasmid (Figure S6e). While the effect is weak in Model II, it is strong in Model III since in that case, due to segregational loss, many plasmid-free cells are already present to receive the rescue plasmid when the environment changes. Nonetheless, if the probability of segregational loss is too high, the 'plasmid-free equilibrium' is the only possible equilibrium. In that case, the only effect of segregational loss is to reduce the effective fitness of rescue plasmid cells, leading to a decreased rescue probability (dashed line in Figure S7b).

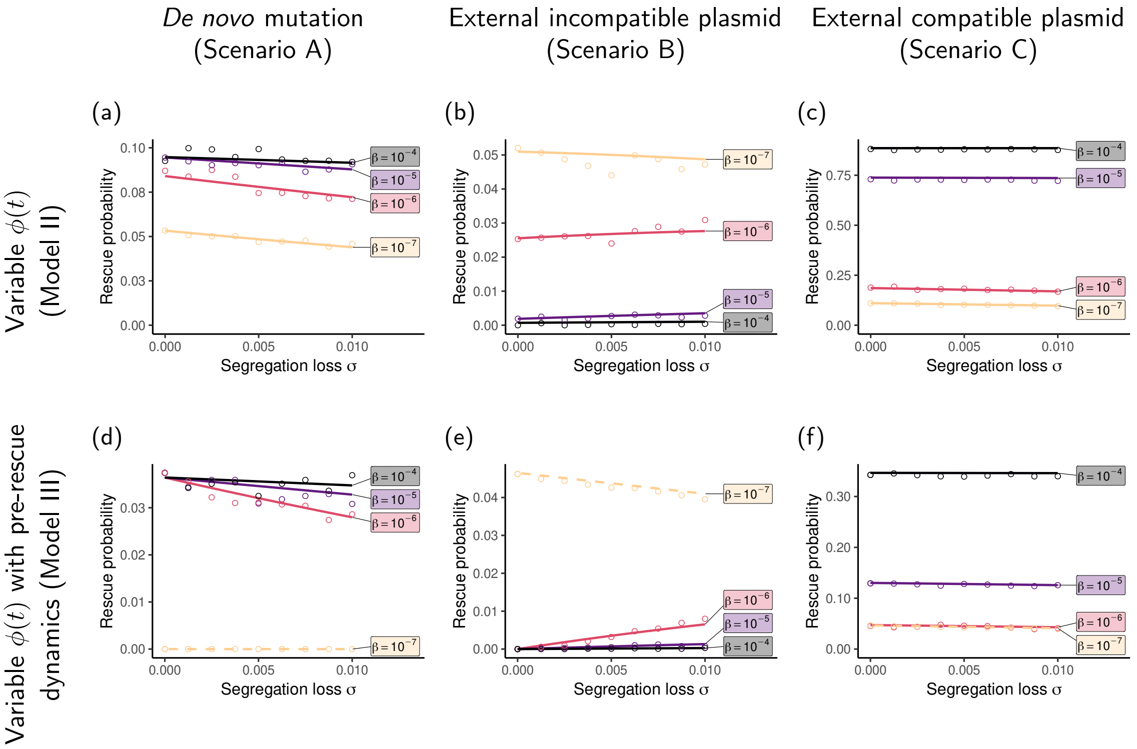


Figure S6: The probability of evolutionary rescue when there is a probability of segregational loss of plasmids ($\sigma$) at cell division. We here assume, as in the main text, that $\beta_r = \beta_R = \beta$. In Panels (a)-(c), the initial fraction of resident plasmid cells is $\phi_0 = 1/2$ and plasmids impose no cost on their hosts ($c = 0$). In Panels (d)-(f), the plasmid cost is $c = 0.01$. All other parameters are the same as in Figures 3, 5, and 6 in the main text. Solid lines indicate values of $\beta$ for which the population consists of a mix of plasmid-free and resident plasmid prior to the environmental change, while the dashed line indicates a value of $\beta$ for which the population consists of only plasmid free cells (low $\beta$). Note that the range of the $y$-axis varies across panels.

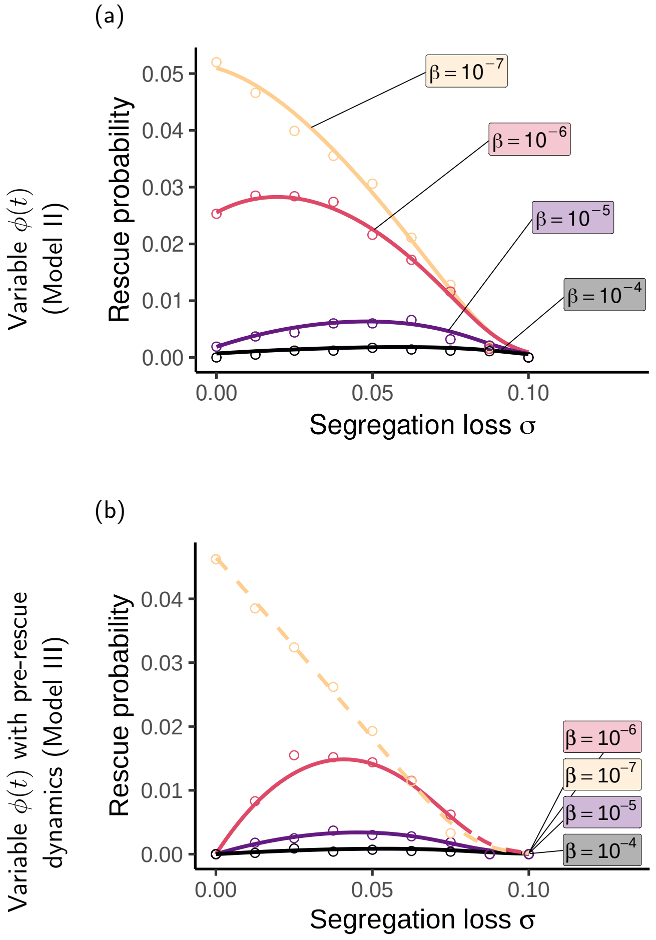


Figure S7: The probability of evolutionary rescue when there is a probability of segregational loss of plasmids ($\sigma$) at cell division. The figure corresponds to Figures S6b and S6e but for higher values of $\sigma$. In Panel (b), solid lines indicate values of $\beta$ for which the population consists of a mix of plasmid-free and resident plasmid cells prior to the environmental change, while dashed lines indicate values of $\beta$ for which the population consists of only plasmid free cells (low $\beta$) or only resident plasmid cells (high $\beta$).

#### S5 Comparison to rescue on the chromosome

In this section, we compare the probabilities of rescue by *de novo* mutations on the chromosome and on a plasmid. For rescue on the chromosome, we consider a resident population of $N(t)$ plasmid-free cells. Its dynamics is given by equation (2) in the main text. Rescue alleles appear in these cells through *de novo* mutation at rate $\lambda(t) = u(1 + s_0)N(t)$. Note that we assume that the mutation rate $u$ is the same for the chromosome and the plasmid. Cells with the rescue allele are modelled by a birth and death process with the following rates:

$$b(t) = 1 + s_R,$$
$$d(t) = 1 + \alpha N(t).$$

Figure S8 compares the probabilities of evolutionary rescue on the chromosome (dashed red line) and on a conjugative plasmid with a variable fraction of resident plasmid cells $\phi(t)$ (Model II). The comparison is meaningful only in the case when rescue genes appear via *de novo* mutation (Scenario A). For many different values of the parameters of our model, rescue is never more likely to occur on the plasmid than on the chromosome. It is usually even more likely to occur on the chromosome. Only when the transfer coefficient $\beta$ is large enough, the probability of rescue on a plasmid converges to the one on the chromosome. This is because, if plasmid-free cells are converted fast enough to resident plasmid cells, then the rate of appearance of adapted cells and the birth-death process are identical for a rescue mutation occurring on the chromosome and on a plasmid.

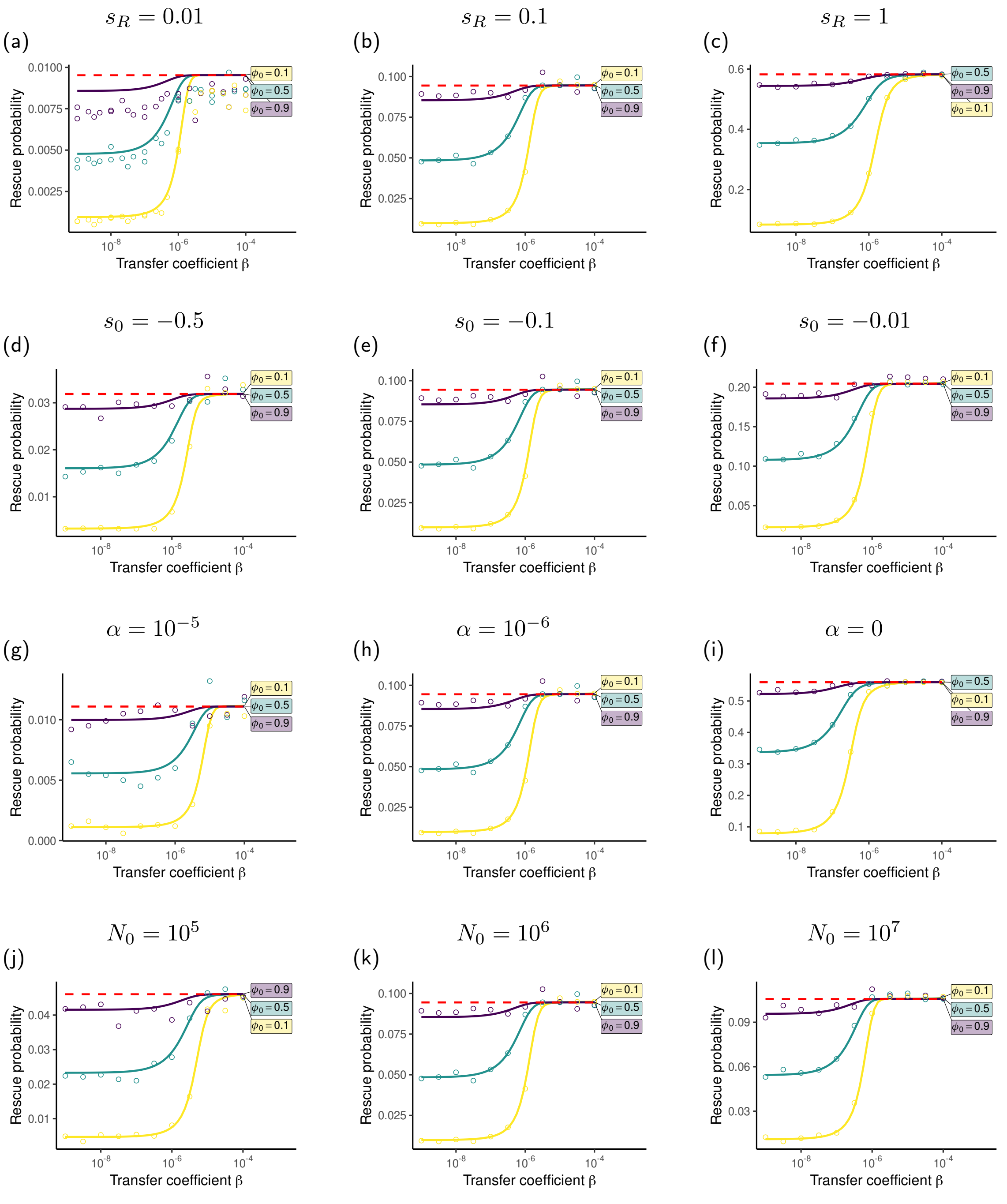


Figure S8: Comparison of rescue via *de novo* mutation on the chromosome (dashed red line) and on a conjugative plasmid with a variable fraction of resident plasmid cells $\phi(t)$ (Model II). Unless stated otherwise above each panel, parameters used are the default values defined in the main text.

#### S6 Additional figures

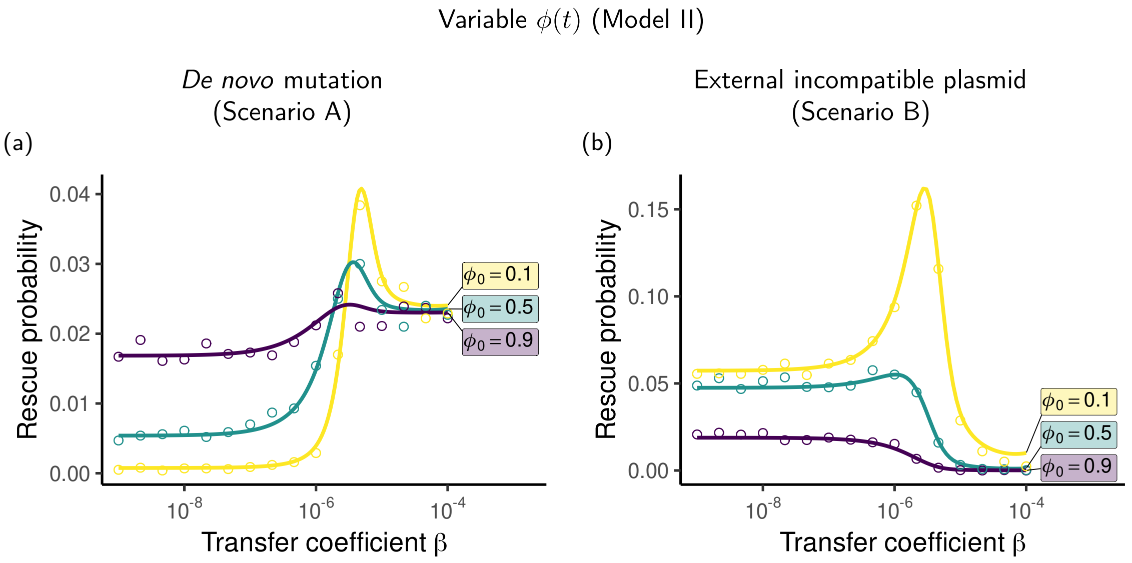


Figure S9: Probability of evolutionary rescue with variable fraction of resident plasmid cells $\phi(t)$ and high plasmid cost $c$. In Panel (a), rescue plasmids appear through *de novo* mutations. In Panel (b), incompatible rescue plasmids appear through transfers from an external population. Parameters used are the default values defined in the main text except for $s_R = 1$ and $c = 1$.
